# Real-time imaging of mitochondrial redox reveals increased mitochondrial oxidative stress associated with amyloid β aggregates *in vivo* in a mouse model of Alzheimer’s disease

**DOI:** 10.1101/2022.06.12.495840

**Authors:** Maria Calvo-Rodriguez, Elizabeth K. Kharitonova, Austin C. Snyder, Steven S. Hou, Maria Virtudes Sanchez-Mico, Sudeshna Das, Zhanyun Fan, Hamid Shirani, K. Peter R. Nilsson, Alberto Serrano-Pozo, Brian J. Bacskai

**Affiliations:** Department of Neurology, Massachusetts General Hospital and Harvard Medical School, 114, 16^th^ St, Charlestown, MA 02129, US; Department of Physics, Chemistry and Biology, Linköping University 581 83 Linköping, Sweden

**Keywords:** Oxidative stress, ROS, Alzheimer’s disease, mitochondria, SS31, neurodegeneration, multiphoton microscopy

## Abstract

**Background:** Reactive oxidative stress is a critical player in the amyloid beta (Aβ) toxicity that contributes to neurodegeneration in Alzheimer’s disease (AD). Mitochondrial damage, observed in AD, is one of the main sources of reactive oxygen species. Although Aβ causes neuronal mitochondria-associated reactive oxidative stress *in vitro*, this has never been directly observed in the *in vivo* living brain. Here, we tested whether Aβ plaques and soluble oligomers induce mitochondrial oxidative stress in surrounding neurons *in vivo*, and whether the neurotoxic effect can be abrogated using mitochondrial-targeted antioxidants.

**Methods:** We expressed a genetically encoded fluorescent ratiometric mitochondria-targeted reporter of oxidative stress in mouse models of the disease, and performed intravital multiphoton microscopy of neuronal mitochondria and Aβ plaques.

**Results:** For the first time, we demonstrated by direct observation exacerbated mitochondrial oxidative stress in neurons after both Aβ plaque deposition and direct application of soluble oligomeric Aβ onto the brain, and determined the most likely pathological sequence of events leading to oxidative stress *in vivo*. Oxidative stress could be inhibited by both blocking calcium influx into mitochondria and treating with the mitochondria-targeted antioxidant SS31.

**Conclusions:** Considering these results, mitochondria-targeted compounds hold promise as neuroprotective drugs for the prevention and/or treatment of AD.

## Introduction

Alzheimer’s disease (AD) is a progressive, neurodegenerative disorder that leads to dementia. The main neuropathological hallmarks of AD are the deposition of extracellular amyloid beta (Aβ) plaques and intraneuronal neurofibrillary tangles, as well as multiple cellular changes, including neuronal and synapse loss, synaptic dysfunction, mitochondrial structural and functional abnormalities, and inflammatory responses. The FDA has recently approved an anti-Aβ antibody as the first therapy targeting the fundamental pathophysiology of the disease. Unfortunately, there is still an urgent necessity of drugs that can prevent or slow AD clinical progression.

Mitochondrial dysfunction is considered an early event in the AD pathophysiologic process. Besides providing ATP, mitochondria play several key roles, including synaptic maintenance, intracellular calcium (Ca^2+^) signaling regulation, free radical production and scavenging, and activation of caspases. An immediate consequence of mitochondria dysfunction is the increase of reactive oxygen species (ROS) production, a byproduct of the electron transport chain, that promotes oxidative damage to DNA, RNA, proteins, and lipids. Mitochondria are both one of the main cellular sources of ROS, and one of the targets of ROS toxicity. Oxidative stress results from an imbalance between pro-oxidants and antioxidants. Neurons hamper mitochondrial oxidative stress with cellular antioxidant defenses and even with removal of damaged mitochondria via mitophagy when there is more severe damage (1). These systems regulate the cellular reduction/oxidation (redox) balance, and thereby, neuronal survival. However, when mitochondria are severely damaged, the antioxidant defense decreases, thus increasing the production of ROS (which further damages mitochondria), enhancing free radicals’ generation, and reducing or depleting the antioxidant capacity.

Oxidative damage and mitochondrial degeneration are involved in AD pathophysiology. For instance, the level of antioxidant enzymes is decreased in plasma from AD patients, thus leading to an accumulation of oxidative damage (2). AD pathogenesis is correlated with increased oxidative stress levels due to enhanced production of ROS and/or decreased antioxidant defense mechanisms (3). However, whether oxidative stress is the cause or consequence of AD neuropathological changes and whether changes in the redox homeostasis have a direct impact on the progression of AD pathology remain open questions. Recently, mitochondria-targeted molecules have been developed and tested as neuroprotective drugs for AD (4, 5). However, no studies have directly visualized the effects of Aβ aggregates on mitochondrial redox state *in vivo*, and whether this can be abrogated using mitochondria-targeted antioxidants.

To address these questions, we used a combination of a genetically encoded fluorescent redox sensor and multiphoton microscopy to image and monitor *in vivo* the mitochondrial redox imbalance in neurons from the APP/PS1 transgenic (Tg) mouse model of AD. We demonstrate that Aβ plaque deposition, and particularly soluble oligomeric Aβ species, lead to increased mitochondrial oxidative stress levels within neuronal mitochondria. We also show that inhibiting mitochondrial Ca^2+^ influx, which is pathologically exacerbated in these Aβ plaque-depositing mice (6) can prevent this mitochondrial redox dysregulation in neurons. Lastly, we demonstrate that the use of antioxidants directly targeting mitochondria can decrease mitochondrial oxidative stress levels to basal levels in the AD mice, supporting the idea that mitochondria-targeted compounds that prevent or minimize mitochondrial dysfunction could hold promise as neuroprotective drugs against AD progression.

## Materials and methods

### 1. Animals

Animal experiments were performed under the guidelines of the Institutional Animal Care and Use Committee (IACUC, protocol #2018N000131). All experimental procedures were approved by the Institutional Animal Care and Use Committee at Massachusetts General Hospital. The following transgenic lines were used: APPswe/PSEN1ΔE9 double Tg mice (The Jackson laboratory, B6.Cg-Tg(APPswe, PSEN1dE9)85Dbo/Mmjax, MMRRC Cat# 034832-JAX, RRID:MMRRC_034832-JAX) (APP/PS1, 2- to 3-months of age and 8- to 10-months of age) of either sex, expressing both human *APP* gene carrying the Swedish mutation and exon 9 deletion mutation in the *PS1* gene, and age-matched non-transgenic littermates (Wt) were used as controls; and C57BL/6J males (4- to 5-months of age, Charles River) for the application of DTT and DTDP, conditioned media and Ru360. For conditioned media preparation, Tg2576 males (Taconic Farms, B6;SJL-Tg(APPSWE)2576Kha, IMSR Cat# TAC:1349, RRID:IMSR_TAC:1349), which heterozygously overexpress human APPswe under the PrP promoter, were mated with Wt females for preparation of primary cortical neurons. Mice were socially housed with up to four animals per cage, with ad libitum access to food and water, in a 12-hour light/dark cycle and controlled temperature and humidity conditions. A sample size of at least 3 mice (of either sex) was randomly allocated to experimental groups.

### 2. Cell culture

Mouse neuroblastoma cells (N2a) were grown at 37C in a humidified incubator chamber under 5% CO_2_ in OptiMEM (Gibco), supplemented with 5% fetal bovine serum (FBS) (Atlanta Biologicals) and 1% penicillin and 1% streptomycin (Gibco) in a humidified 37 C incubator with 5% CO_2_. Cells were plated into 8-well chamber (Sarstedt) at a density of 30,000 cells/well and transiently transfected 24h later using Lipofectamine 2000 (Life Technologies) according to the manufacturer’s instructions and imaged 24h after.

Primary cortical neurons were prepared as previously described (6). Briefly, neurons were obtained from embryonic day 14 (E14) CD1 (Charles Rives Laboratories) mouse embryos. Neurons were dissociated using Papain dissociation system (Worthington Biochemical Corporation, Lakewood, NJ, USA). Cells were plated in 8-well chamber slides previously coated with poly-D-lysine at a density of 30,000 cells/well and were maintained for 10-14 days *in vitro* (DIV) in Neurobasal medium supplemented with 2% B27 (Gibco), 1% penicillin/streptomycin (Gibco) and 1% glutamax (Gibco) in a humidified 37 C incubator with 5% CO_2_ without further media exchange. Neurons were transfected using Lipofectamine 2000 (Life Technologies) or infected with AAV.hSyn.mt-roGFP after 12-14 DIV, and imaged 1 or 3 days later respectively. Experiments were performed after a culturing period of 12-14 DIV.

### 3. Plasmids

mRuby-Mito-7 was a gift from Michael Davidson (Addgene plasmid #55874; http://n2t.net/addgene:55874; RRID:Addgene_55874). mRuby-ER-5 was a gift from Michael Davidson (Addgene plasmid #55860; http://n2t.net/addgene:55860; RRID:Addgene_55860). Matrix-roGFP (mt-roGFP) was a gift from Paul Schumacker (Addgene plasmid #49437; http://n2t.net/addgene:49437; RRID:Addgene_49437).

### 4. AAV.hsyn.mt-roGFP construction and production

DNA sequence of mt-roGFP (8) was ligated between the inverted terminal repeat sites (ITRs) of an adenovirus (AAV) packaging plasmid with human synapsing (hSyn) promoter, and WPRE/SV40 sequence. The expression cassette included the following components: (1) a 1.7-kb sequence containing human synapsin 1 gene promoter, (2) mt-roGFP, (3) WPRE, and (4) Simian virus 40 (SV40). Human embryonic kidney (HEK) 293T cells were co-transfected with the construct and a helper plasmid and harvested. Virus was purified and titrated by infecting HEK293T cells. Virus titer was 1.0 x 10^12^ viral genome copies per mL.

### 5. SS31 preparation and drug delivery

SS31 (D-Arg-Dmt-Lys-Phe-NH_2_; Dmt = 2’,6’-dimethylthyrosine) and SS20 (Phe-D-Arg-Phe-Lys-NH_2_) were obtained from Biomatik (https://www.biomatik.com). SS31 and SS20 were administered intraperitoneally to APP/PS1 Tg and non-transgenic littermate mice (5mg/kg body weight) twice a week for 8 weeks. The treatment began when the mice were 8 months of age, and the recording sessions were carried out at 10 months of age. All mice were daily observed by a veterinarian. SS31 dose was determined based on previous studies (37, 72), suggesting that this concentration had the maximum protective effects without any side-effects or toxicity.

### 6. Preparation of Wt and Tg neuronal conditioned media and Aβ-immunodepleted media, and measurement of soluble Aβ levels

Primary cultures from Tg2576 mice, heterozygous for the hAPPswe mutation, were prepared as explained above. Tissue from each individual embryo was collected for genotyping. Primary neurons were maintained in Neurobasal media containing 2% B27 supplement, 1% penicillin/streptomycin and 1% Glutamax. Conditioned media from either Tg cultures (TgCM) or Wt littermates (WtCM) was collected at 14 DIV. Measurement of soluble Aβ levels were conducted using sandwich ELISA as previously described (6). Briefly, CM was collected from the primary cultures, and Aβ_1-40_ was measured with commercial colorimetric ELISA kit (WAKO, Wako #294-64701 Human/Rat Aβ_1-40_), specific for human. A 96-well plate reader was used, following the manufacturer’s instructions. Each sample was run in duplicates. Protein concentrations of the CM were determined and Aβ was expressed in nM. Additionally, human Aβ was immunodepleted from TgCM with the mouse antibody 6E10 (Purified anti-β-Amyloid, 1-16 antibody BioLegend Cat# 803004, RRID: AB_2715854) and Protein G Sepharose beads (Sigma-Aldrich). Protein G beads were conditioned with cold Neurobasal media. 1 ml of TgCM and 40 μl of the pre-conditioned G beads were incubated with 6 μg of 6E10 antibody overnight at 4C under rotation. Supernatant was collected and Aβ concentration quantified by colorimetric ELISA human/rat Aβ_1-40_. Aβ_1-40_ concentration in the media was WtCM 0.3 nM, TgCM 5 nM and Aβ-immunodepleted TgCM 0.7 nM. The concentration of Aβo in the TgCM has been shown to represent around 10% of the total amount of Aβ_1-40_(20).

### 7. Stereotactic intracortical injection of AAV.hsyn.mt-roGFP

For acute experiments, AAV.hSyn.mt-roGFP was injected into four- to five-month old C57BL/J6 Wt mice somatosensory cortex, as previously described (7). Mice were anesthetized using 5% isoflurane (vol/vol) for induction and maintained at 1.5% throughout the surgery. Under a stereotactic frame (Kopf Instruments), a burr hole was drilled in the skull at 1mm anteroposterior, 1 mm mediolateral from bregma, and AAV was injected at −0.8 mm dorsoventral. A programmable syringe pump with a 33-gauge sharp needle attached to a 10 μL Hamilton micro-syringe was used for infusion. 3 μL of viral suspension were injected at 0.15 μL/min. Body temperature was maintained throughout surgery with a heating pad. After injection, the needle was removed, and the mouse scalp sutured. Mice were put on a heating pad for recovery. For stable fluorescent expression, mice were imaged 3 weeks after injection. For chronic experiments, AAV.hSyn.mt-roGFP was injected to 9 mo-old or 3 mo-old Tg mice in the somatosensory cortex at the moment of the cranial window implantation. Mice were given buprenorphine (0.1mg/kg) for 3 days following surgery.

### 8. Cranial window implantation

Mice were anesthetized with isoflurane, the scalp shaved and sterilized, and an incision was made to expose the underlying skull. A custom-made stereotax fixed the skull. For chronic windows, an area of skull no larger than 5-mm was removed and replaced with a glass coverslip for imaging. After craniotomy, mice were put on a heating pad for recovery and two-photon imaging was performed 3 weeks later. For acute windows (*in vivo* validation and CM experiments), dura matter was removed, and 6 mm windows were implanted. Two-photon imaging was performed after cranial window implantation. Mice were given buprenorphine (0.1mg/kg) for 3 days following surgery.

### 9. In vivo multiphoton microscopy imaging

HS169 (10 mg/kg) was retro-orbitally injected 24h before the imaging session to label amyloid plaques (17). Texas Red dextran (70,000 MW; 12.5 mg/mL in PBS; Molecular Probes) was retro-orbitally injected to provide a fluorescent angiogram. Mice were anesthetized by isoflurane and head-restrained using a custom made stereotax. Images of AAV.hSyn.mt-roGFP were acquired on an Olympus FluoView FV 1000MPE multiphoton laser-scanning system mounted on an Olympues BX61WI microscope and equipped with a 25x Olympus water immersion objective (1.05 numerical aperture (NA)). A Deep-See Mai Tai Ti:Sapphire mode-locked laser (Spectra-Physics) was used for multiphoton excitation at the following wavelengths: 800 and 900 nm for AAV.hSyn.mt-roGFP, 800 nm for HS169 and 900 nm for Texas Red Dextran. Emitted fluorescence was collected in three channels in the range of 460-500, 520-560 and 575-630 nm. All images were obtained at depths up to 200 μm from the pial surface. All images were captured at a 5x digital zoom. Five to eight cortical volumes (Z-series, 127 µm x 127 µm) were acquired per mouse, at a step size of 2 µm and a resolution of 512 x 512 pixels. Photomultiplier settings remained unchanged throughout the different imaging sessions. Laser power was adjusted as needed to avoid image saturation and kept below 30 mW to avoid phototoxicity.

For acute CM experiments and *in vivo* validation of AAV.hSyn.mt-roGFP, an imaging session was first performed to determine the basal resting ratio 800/900. Then, the window was opened and sealed again with DTT, DTDP, WtCM, TgCM, Aβ-immunodepleted TgCM, Ru360 (Calbiochem, Merck Millipore) or Ru360 + TgCM (40 μL final volume applied). Ru360 was preincubated for 15 min previous to application of TgCM. After either 20 minutes (for validation) or 1h (for CM experiments), the same fields of view were reimaged to determine the relative changes in ratio 800/900 (ΔR/R_0_). The fluorescent angiogram created by Texas Red Dextran helped with re-locating the same fields of view.

### 10. Image processing and quantification

RoGFP displays two excitation peaks that are sensitive to redox changes. The redox status is assessed by monitoring the ratio of GFP fluorescence emission at 800 and 900 nm excitation (43, 75). For the analysis of mitochondrial redox state in primary neurons, ROIs were drawn around somas and primary processes, and in background regions outside cells. Pixel intensity was measured on images taken at two wavelengths (800 and 900 nm), and background subtracted. To calculate the fluorescent intensity ratios, the 800 nm image was divided by the 900 nm image in a pixel-by-pixel manner.

All multi-photon images were analyzed using customized MATLAB scripts (MathWorks). To perform automatic segmentation of mitochondria, an adaptive thresholding procedure was applied to the *in vivo* images (acquired at 5x digital zoom) to generate binary images and individual mitochondria in neuronal somas and processes were identified using a constraint on object size. Amyloid plaques were manually extracted from the z-stacks. Background was subtracted from the 800 and 900 nm channel images. The ratio value of each segmented mitochondria was determined by taking of the sum of the pixels within the segmented region in the 800 nm channel and dividing by the sum in the 900 nm channel. Pseudocolored images were generated using MATLAB by first creating the ratio image by dividing the 800 nm image by the 900 nm image on a pixel-by-pixel basis. Next, the ratio image was assigned to the RGB (Red, Green, Blue) colorspace with the color range determined by the maximal achievable redox changes that can be accomplished with DTT and DTDP treatment *in vivo*. The RGB image was then converted to the HSV (Hue, Saturation, Value) colorspace with the Value field being set to the overall intensity image (800 nm + 900 nm channels). Distance to plaque was measured based on the centroid of the mitochondrion to the edge of the nearest plaque. Images presented in the figures are a single slice from the z-stack.

### 11. Immunohistochemistry

Animal subjects were euthanized and perfused with 20 mL of PBS followed by 20 mL of 4% PFA. Brains were extracted and kept in a fixing solution (4% PFA and 30% glycerol in PBS) for 24h, and then embedded in OCT. The OCT block was then sectioned into 20 µm coronal sections on a cryostat (Leica). Sections were subjected to antigen retrieval first, by heating with citrate buffer with Tween20 at pH 6.0, and then permeabilized with 0.5% triton X-100, blocked with 5% normal goat serum, and incubated with target antibodies at 4C o/n (GFP (1:500, Antibodies Incorporated Cat# GFP-1020, RRID:AB_10000240), HSP60 (1:200, Abcam Cat# ab46798, RRID:AB_881444), GS (1:500, Abcam Cat# ab73593, RRID:AB_2247588), NeuN (1:500, R&D Systems Cat# MAB377, RRID:AB_2298767), Amyloid β (1:500, Immuno-Biological Laboratories Cat# 18584, RRID:AB_10705431), Neurofilament SMI312F (1:1000, BioLegend Cat# 837904, RRID:AB_2566782). Corresponding secondary antibodies (Alexa Fluor 488 1:1000 (Molecular Probes Cat# A-11039, RRID:AB_142924), Alexa Fluor 647 1:1000 (Molecular Probes Cat# A-21245, RRID:AB_141775 and Molecular Probes Cat# A-21235, RRID:AB_2535804), Alexa 594 1:1000 (Molecular Probes Cat# A-11012, RRID:AB_141359), Cy3 1:1000 (Molecular Probes Cat# A-11039, RRID:AB_142924 and Abcam Cat# ab6939, RRID:AB_955021), Cy5 1:1000 (Abcam Cat# ab97035, RRID:AB_10680176 and Jackson ImmunoResearch Labs Cat# 111-175-144, RRID:AB_2338013) were applied and incubated for 1h at room temperature. Appropriate sections were treated with 1% ThioS, and/or cover slipped with antifade mounting medium with DAPI (Vector Laboratories, (H1500)).

### 12. Confocal microscopy

Imaging of sections ((GFP, HSP60, NeuN, GS), and (Aβ, NF)) was conducted on an Olympus Fluoview 3000 confocal microscope with a 40x immersion objective. Applicable laser excitation wavelengths were used for optimal fluorophore excitation and emission separation and detection. Parameters were maintained for all slides within one antibody condition.

### 13. Fluorescence imaging

For plaque burden analysis (Aβ immunohistochemistry and ThioS), slides were scanned using an Olympus VS120-S6-W virtual slide microscope. Images were taken with a 20x objective. Acquired images were analyzed using CellSens software (Olympus). A manual threshold was set to include the diffuse and dense plaque cores. These parameters were maintained throughout all image analyses and analyzed for total burden.

### 14. Analysis of publicly available human brain single RNA-seq datasets

The expression of 31 mitochondrial genes involved in mitochondrial antioxidant defense was compared in advanced AD Braak stages (V-VI) versus control subjects (Braak 0-I-II). Data was obtained from a single-nucleus RNA-Seq performed on prefrontal cortex samples from 48 individuals with varying levels of AD pathology (46). The samples were grouped by Braak score, which assesses the distribution of tau neurofibrillary tangles in the subject’s brain (76). Multiple comparisons corrections were performed using the Benjamini-Hochberg method (77). For the purpose of selection for visual display, an FDR threshold of 0.25 was used.

### 15. Statistics

Graph Pad Prism (version 6.0) was used for statistical analysis and data presentation. Data are reported as mean ± SEM. Mann-Whitney t test was used to compare two different conditions. Kruskal-Wallis test and Dunn’ multiple comparisons test were used to compare different conditions (i.e. CM). Statistical correlations were determined using Pearson’s correlation coefficient. In each experiment, the number of animals, volumes, and statistical parameters can be found in the figure legend. Resulting mean and standard error estimates and the p-values are presented in the Figure legends. p<0.05 was considered statistically significant.

## Results

### 1. hsyn.mt-roGFP targets mitochondria and reports oxidative stress changes in vitro and in vivo

To study specific redox changes, we used the genetically encoded redox-sensitive fluorescent protein roGFP (7) targeted to the mitochondrial matrix (matrix-roGFP (8), heretofore mt-roGFP) by using the mitochondria-targeting sequence from cytochrome oxidase subunit IV (8). The ratio of the fluorescence after excitation at 800 nm over the excitation at 900 nm (Ratio 800/900) reflects the redox environment, with an increase in ratio indicating oxidation. We chose this reporter because of its ratiometric properties (ratiometric by sequential excitation near its absorption maxima at 800 and 900 nm), which enables true quantitative redox imaging (9, 10). RoGFP redox sensors almost exclusively report reduced/oxidized glutathione (GSH/GSSG) ratio (11), which is mediated by cell endogenous glutaredoxins. Additionally, roGFP has been described to be minimally affected by cellular pH and Cl^-^ changes in the physiological range (12, 13), unlike other ratiometric redox sensors such as Hyper (14), which senses H_2_O_2_ but markedly responds to pH changes (15). Mt-roGFP is restricted to the mitochondrial matrix and is not associated with either cristae or peripheral mitochondrial membranes ((8) and Supplementary Figure 1).

We first validated the sensor specificity in cultured N2a cells and primary mouse neurons by co-labeling with a different plasmid targeted to mitochondria. Colocalization of mt-roGFP and mRuby-mito7 confirmed the mitochondrial localization of mt-roGFP (Supplementary Figure 1a). In addition, lack of colocalization between mt-roGFP and an endoplasmic reticulum (ER) marker (mRuby-ER5), co-transfected in N2a cells, further demonstrated the mitochondria-specific targeting of mt-roGFP (Supplementary Figure 1b).

We then validated the sensor functionality in cultured primary mouse neurons by determining its responsiveness to reduction/oxidation. Primary neurons were exposed to saturating doses of the reductant DTT (dithiothreitol) or the oxidant DTDP (dithiodipyridine), to fully reduce or oxidize the cells respectively. As illustrated in Supplementary Figure 1c and d, treatment with DTT resulted in an expected increase in fluorescence intensity at 900 nm (in green), and a decrease at 800 nm (in red), and vice versa under the oxidizing conditions of DTDP treatment. The Ratio 800/900 distribution was slightly shifted to the left under DTT reducing conditions, indicating lower redox levels in the mitochondrial matrix (Supplementary figure 1d blue histogram), and greatly shifted to the right under DTDP oxidizing conditions (Supplementary Figure 1d red histogram), indicating higher relative oxidized levels.

We next targeted the reporter to neurons by using the human synapsin promoter (hSyn), and packaged it into an adenoviral-associated vector (AAV.hSyn.mt-roGFP) (Figure 1a). We validated its localization by injecting it in the somatosensory cortex and co-staining with markers for neurons (NeuN, neuronal nuclei antigen, Fox3), astrocytes (GS, glutamine synthetase) and mitochondria (HSP60, heat shock protein 60). Supplementary Figure 2 shows the exclusive localization of the vector to neuronal mitochondria (but not astrocytes), based on its colocalization with NeuN and HSP60. We then tested whether it was possible to monitor dynamic changes in mitochondrial oxidative stress in neurons *in vivo* with the newly created AAV.hSyn.mt-roGFP reporter. Brains of wild-type (Wt) living mice were topically exposed to the reductant DTT or the oxidant DTDP through a craniotomy, to fully reduce or oxidize their neuronal mitochondria respectively, and their effects were imaged with multiphoton microscopy. Treatment with DTDP increased the relative oxidative stress levels within the mitochondria of neurons (represented as Ratio 800/900), whereas DTT did not alter the relative redox levels as measured by the sensor (Figure 1b-d). The corresponding histograms show a shift toward higher ratios upon hSyn.mt-roGFP oxidation with DTDP, but no change upon reduction with DTT (Figure 1e, f), consistently with the greater effect of DTDP over DTT observed *in vitro* (Supplementary Figure 1d).

**Figure 1.**
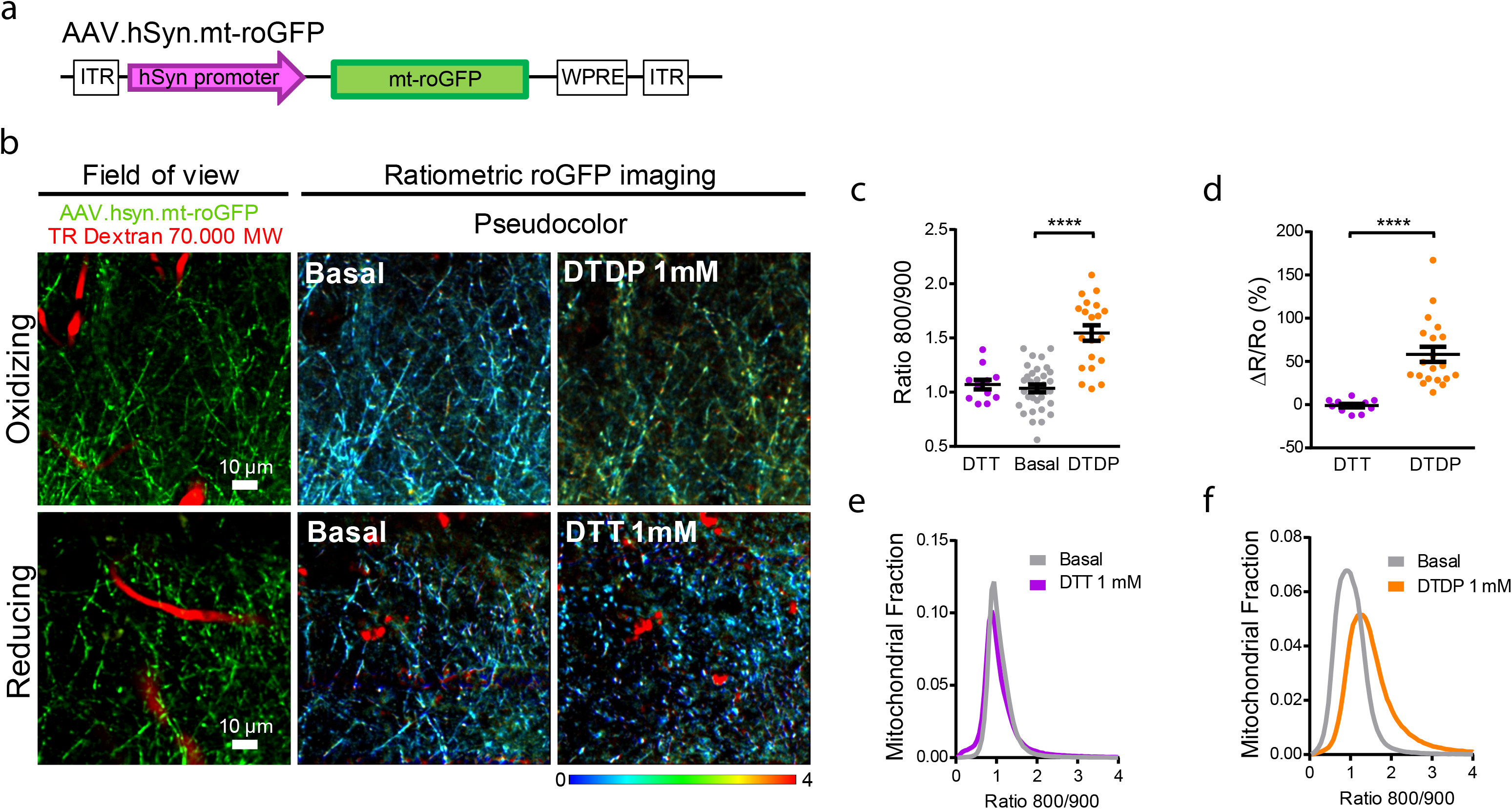
AAV.hSyn.mt-roGFP expresses in neuronal mitochondria and is functional *in vivo.* **a.** Diagram of construct of AAV.hSyn.mt-roGFP.WPRE. **b.** Validation of pAAV.hSyn.mt-roGFP *in vivo.* 4-month-old C57Bl/6 mice were intracortically injected with pAAV.hSyn.mt-roGFP and exposed to either the oxidant DTDP or the reducing agent DTT. Field of view shows two-photon images of the expression of the AAV (green) in the cortex of the C57BL/6J mice. Blood vessels are labelled with Texas Red (TR) Dextran 70,000 MW (red). Ratiometric roGFP imaging shows pseudocolor images according to the pseudocolor scale on the bottom, in basal conditions and after topical application of DTDP or DTT onto the brain surface. Scale bars represent 10 μm. **c**. Scatter dot plot represents the Ratio 800/900 in neuronal mitochondria of every volume acquired. Error bars represent mean ± SEM. (Basal: 1.06 ± 0.036, DTT: 1.07 ± 0.044, DTDP: 1.55 ± 0.072; ****p<0.0001, n = 33, 12 and 20 z-stacks from 7, 3 and 4 mice respectively). **d.** Scatter dot plot represents the relative change (%) in ratio for each condition. Error bars represent mean ± SEM (DTT: −0.96 ± 1.997%, n = 12 z-stacks; DTDP: 58.18 ± 8.719%, n = 20 z-stacks from 3 and 4 mice respectively, ****p<0.0001). **e, f.** Histograms of Ratio 800/900 frequency distribution in basal conditions (grey) and 20 min after application of DTT 1 mM (**e**, purple) and DTDP 1 mM (**f**, orange) in neurons.

Taken together, these data demonstrate that hSyn.mt-roGFP sensor enables monitoring of mitochondrial relative oxidized levels of cortical neurons in living mice. We next extended these findings to a mouse model of cerebral β-amyloidosis.

### 2. Mitochondrial oxidative stress in neurons from AD transgenic mice after Aβ plaque deposition

We used the APPswe/PSEN1ΔE9 (APP/PS1) Tg mouse as a model of AD. This mouse model deposits amyloid plaques starting at 5 months of age (16). In order to analyze redox state changes due to Aβ aggregates, we first evaluated mitochondrial redox levels after Aβ plaque deposition. APP/PS1 Tg and non-Tg littermate mice were injected with AAV.hSyn.mt-roGFP followed by cranial window implantation (Figure 2a). HS169 dye was injected IV 24h before the imaging session to label Aβ plaques (17), and Texas red dextran was injected on the same day to create a fluorescent angiogram (Figure 2b). We observed significant changes in the mitochondrial redox balance in neurons in the APP/PS1 Tg mouse compared to non-Tg mice (Figure 2c, d, e). To determine whether Aβ plaques impact mitochondrial oxidative stress levels in the surrounding neurons, a Pearson correlation between the Ratio 800/900 and the distance to the nearest Aβ plaque was performed (Figure 2f). A negative correlation was observed between the two parameters, suggesting that Aβ plaques could trigger this increase in mitochondrial oxidative stress.

**Figure 2.**
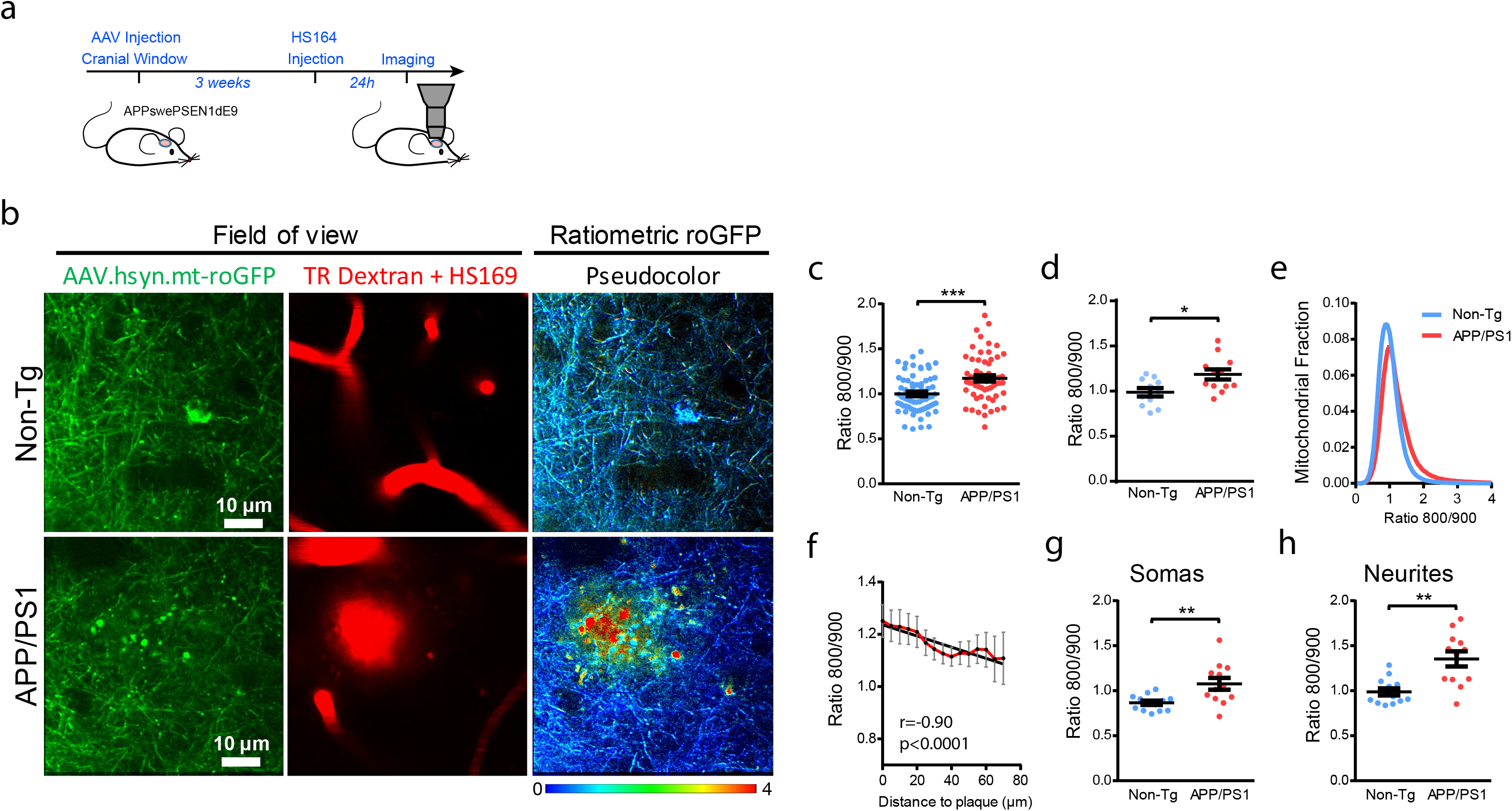
Mitochondrial oxidative stress in AD transgenic mouse neurons after Aβ plaque deposition. **a.** Experimental procedure to determine oxidative stress in neuronal mitochondria in mice. APP/PS1 Tg and non-Tg mice were injected with AAV.hSyn.mt-roGFP and a cranial window was implanted. Three weeks later, oxidative stress was assessed by two photon microscopy. Dextran 70 kDa was used to create a fluorescent angiogram. HS-169 was used to label plaques. **b.** *In vivo* images of neurites and cell bodies expressing AAV.hSyn.mt-roGFP in mitochondria in non-Tg (top) and APP/PS1 Tg mice (bottom). Field of view shows two-photon images of the expression of the AAV (green), blood vessels (Dextran, red) and amyloid plaques (HS169, red) in the cortex. Ratiometric roGFP imaging shows pseudocolor images according to the pseudocolor scale on the bottom. Scale bar represents 10 μm. **c,d.** Mitochondrial oxidative stress (Ratio 800/900) comparison between non-Tg and APP/PS1 Tg mice at 10 months of age, after plaque deposition, in mitochondria in neurons (**c**. average per field of view. non-Tg: 1.00 ± 0.024, n = 69 z-stacks; APP/PS1: 1.17 ± 0.034, n = 60 z-stacks from 11 and 12 mice respectively, ***p = 0.0001. **d**. average per mouse. non-Tg: 0.99 ± 0.046, n = 11 mice; APP/PS1: 1.19 ± 0.056, n = 12 mice, p=0.0224). Error bars represent mean ± SEM. **e**. Histogram of mitochondrial oxidative stress frequency distribution (indicated by Ratio 800/900) in neurons in non-Tg (blue) and APP/PS1 Tg mice (red). **f**. Within 70 µm distance from the edge of a plaque, the probability of finding mitochondrial oxidative stress in neurons was higher close to the plaque (mean ± SEM; *r* (Pearson’s correlation coefficient) −0.90, R square 0.81, ****p<0.0001, n = 33 plaques analysed from 8 Tg mice). **g,h**. Comparison of mitochondrial oxidative stress (Ratio 800/900) in somas and neurites in 10-mo-old non-Tg and APP/PS1 Tg mice. APP/PS1 Tg mice showed significantly higher oxidative stress levels in mitochondria in both compartments. Error bars represent mean ± SEM (**g**. Neuronal somas: 0.87 ± 0.024, n = 13 z-stacks from 4 non-Tg mice, and 1.08 ± 0.067, n = 12 z-stacks from 6 APP/PS1 Tg mice. **p = 0.006. **h**. Neurites: 0.99 ± 0.039, n = 13 z-stacks from 4 non-Tg mice, and 1.35 ± 0.084, n = 12 z-stacks from 6 APP/PS1 Tg mice. **p = 0.002).

Since neurons are highly compartmentalized cells and might have different energy requirements in different compartments (18), we calculated and compared mitochondrial relative redox state in both somas and neurites independently (Figure 2g, h and Supplementary Figure 3a, b). The mitochondrial redox state in neurites was significantly higher than in somas both for APP/PS1 Tg and non-Tg mice (Supplementary Figure 3a, b), with APP/PS1 Tg mice exhibiting higher oxidative stress in both compartments (Figure 2g, h).

To unequivocally determine the role of Aβ as trigger of the observed increased oxidative stress in neuronal mitochondria, we also evaluated mice at 3 months of age, that is, prior to Aβ plaque deposition. The mitochondrial redox state did not significantly differ between young APP/PS1 Tg and non-Tg mice (Supplementary Figure 4). Interestingly, the mitochondrial redox state was also higher in neurites that in somas in both groups of mice at this younger age (Supplementary Figure 3c-d).

Taken together, these data demonstrate that mitochondrial oxidation is increased in the APP/PS1 Tg mice once Aβ plaques have been deposited.

### 3. Soluble Aβ oligomers contribute to the oxidative stress of neuronal mitochondria in vivo

Aβ plaques are surrounded by a halo of soluble Aβ oligomers (Aβo) (19), therefore we aimed to investigate how soluble Aβo directly contribute to the oxidative stress insult in neuronal mitochondria observed in the APP/PS1 Tg mouse. To this end, we used naturally secreted soluble Aβo, previously characterized and shown to contain low molecular weight Aβo (20, 21). Aβ-enriched medium was obtained from primary neurons prepared from Tg2576 embryos (heretofore transgenic conditioned media, TgCM). As control conditions, we used conditioned media from Wt littermates (Wt conditioned media, WtCM) and Aβ-immunodepleted TgCM, which was obtained after exposing TgCM to anti-β-amyloid antibody (6E10, amino acids 1-16). For these experiments, C57Bl/6J mice were injected with AAV.hSyn.mt-roGFP and, after stable expression, the naïve brains were first imaged in baseline conditions (before). Then, either WtCM, TgCM or Aβ-immunodepleted TgCM was topically applied onto the mouse brain, and the same cortical volumes were re-imaged (after), allowing for direct comparation of the Ratio 800/900 between both timepoints (Figure 3a). Representative images of mitochondrial oxidative stress in neurons before and after CM exposure are shown in Figure 3b. We found that the mitochondrial redox state was increased only after application of TgCM, whereas WtCM and Aβ-immunodepleted TgCM (Figure 3c-g) did not alter it. Changes were relatively small, but comparable to the changes observed in the APP/PS1 Tg mouse model. Although only a small fraction of mitochondria is affected by Aβ aggregates, these results support a detrimental role of soluble Aβo around Aβ plaques in the increased mitochondrial redox state *in vivo*.

**Figure 3.**
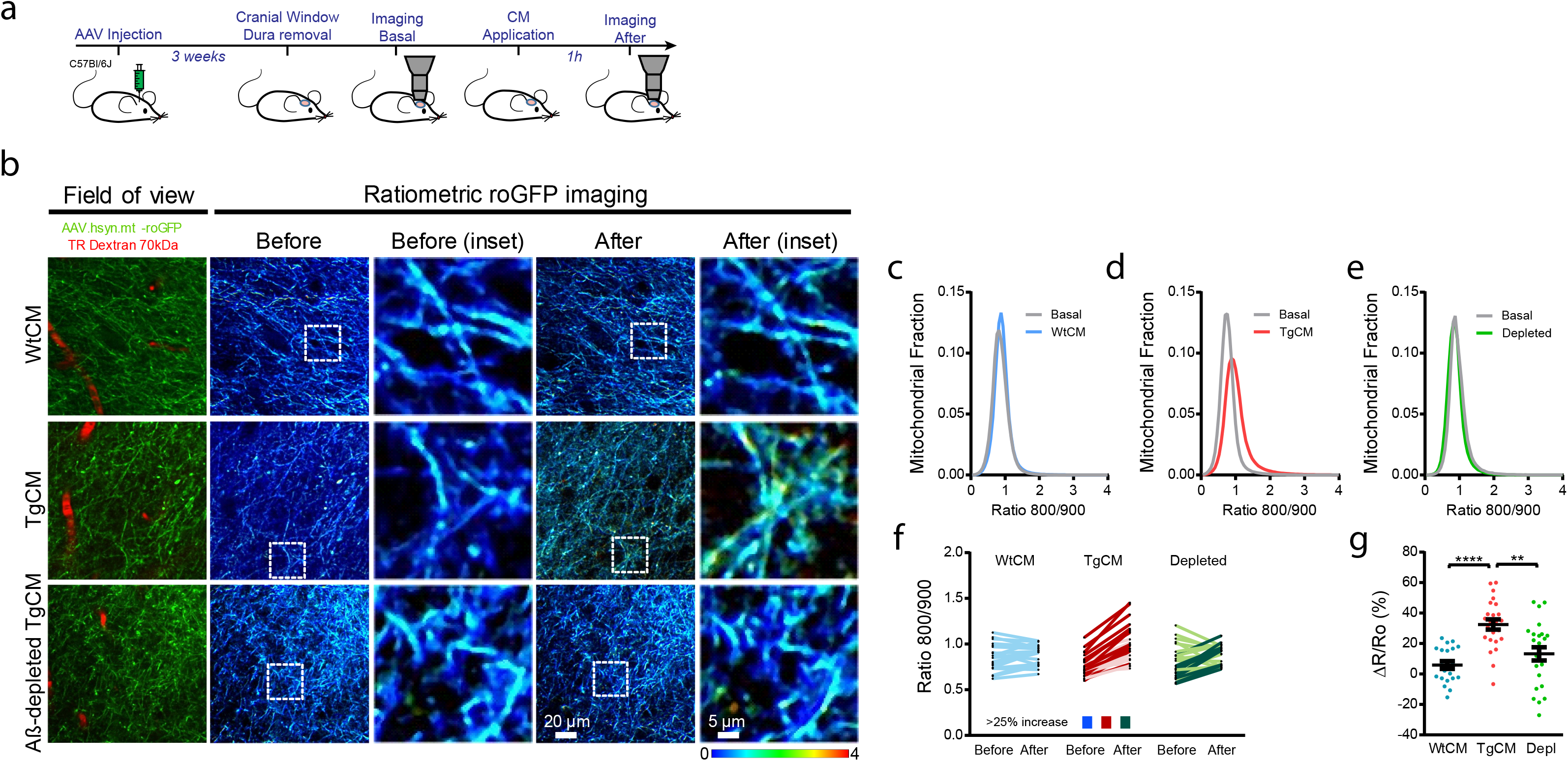
Soluble Aβ oligomers increase oxidative stress in mitochondria in neurons *in vivo.* **a.** Experimental procedure to determine the effects of Aβo on mitochondrial oxidative stress in the healthy mouse brain *in vivo*. 4-months old C57Bl/6 mice were injected with AAV.hSyn.mt-roGFP and a cranial window was implanted 3 weeks later. Oxidative stress in mitochondria was assessed by two photon microscopy in basal conditions and after topical application of either WtCM, TgCM or Aβ-immunodepleted TgCM. **b.** Representative pictures of the effects of CM in neuronal mitochondria in the Wt mouse brain. Scale bar represents 20 μm and 5 μm in the insets. Only TgCM was able to increase oxidative stress levels. **c - e.** Graphs show histograms of mitochondrial oxidative stress frequency distribution (Ratio 800/900) for the three conditions before (basal) and after application of CM in neurons. **f.** Averaged 800/900 ratios before and after CM treatment for each z-stack acquired. Darker traces represent z-stacks showing an increase ≥ 25% in 800/900 ratio after topical application of CM. **g.** Scatter dot plot represents the relative change (ΔR/Ro) in ratio for each condition. Error bars represent mean ± SEM (WtCM: 5.90 ± 2.534%, n=21 z-stacks; TgCM: 32.50 ± 3.328%, 24 z-stacks; depleted TgCM: 13.20 ± 4.340%, 24 z-stacks from 4, 4 and 5 mice respectively, **p<0.01, ****p<0.0001).

### 4. Inhibition of calcium influx into mitochondria prevents neurons from Aβ-induced mitochondrial oxidative stress

An important role of mitochondrial calcium (Ca^2+^) overload in the oxidative stress-induced neuronal cell death and in excitotoxicity has been proposed (22, 23). Mitochondrial Ca^2+^ signaling stimulates oxidative phosphorylation (OXPHOS) and ATP synthesis (24). However, excessive entry of Ca^2+^ into mitochondrial matrix, i.e., mitochondrial Ca^2+^ overload, causes detrimental effects, including mitochondrial membrane potential loss, OXPHOS uncoupling, mitochondrial permeability transition pore opening and eventual cell death (25, 26). Additionally, we have previously demonstrated Aβ-induced mitochondrial Ca^2+^ overload in APP/PS1 Tg mice, as well as after application of soluble Aβo onto the Wt naïve brain (6), and that there is a link between mitochondrial Ca^2+^ overload and the rare neuronal cell death events occurring in these mice (6, 27). Here we investigated whether the Aβ-driven changes in mitochondrial redox levels shown above are mediated via increased mitochondrial Ca^2+^ influx into the mitochondrial matrix. The mitochondrial calcium uniporter (MCU) is the main entrance of Ca^2+^ into mitochondria (28, 29). Following the same experimental protocol as in Figure 3a, we found that blockage of the mitochondrial Ca^2+^ uniporter with the specific cell-permeable MCU inhibitor Ru360 (30) (100 µM) prevented overproduction of mitochondrial ROS, measured as Ratio 800/900, upon application of TgCM onto the Wt mouse brain (Figure 4). Figure 4a shows representative images of mitochondria in baseline conditions (before) and after treatment with TgCM (after) in presence or absence of Ru360, and with Ru360 alone. Since pretreatment with Ru360 abrogated the Aβo-induced increase in mitochondrial oxidation (Figure 4b-f), these findings indicate that increased oxidative stress levels may be downstream of mitochondrial Ca^2+^ overload and that inhibition of MCU exerts a neuroprotective effect on Aβ-induced oxidative stress.

**Figure 4.**
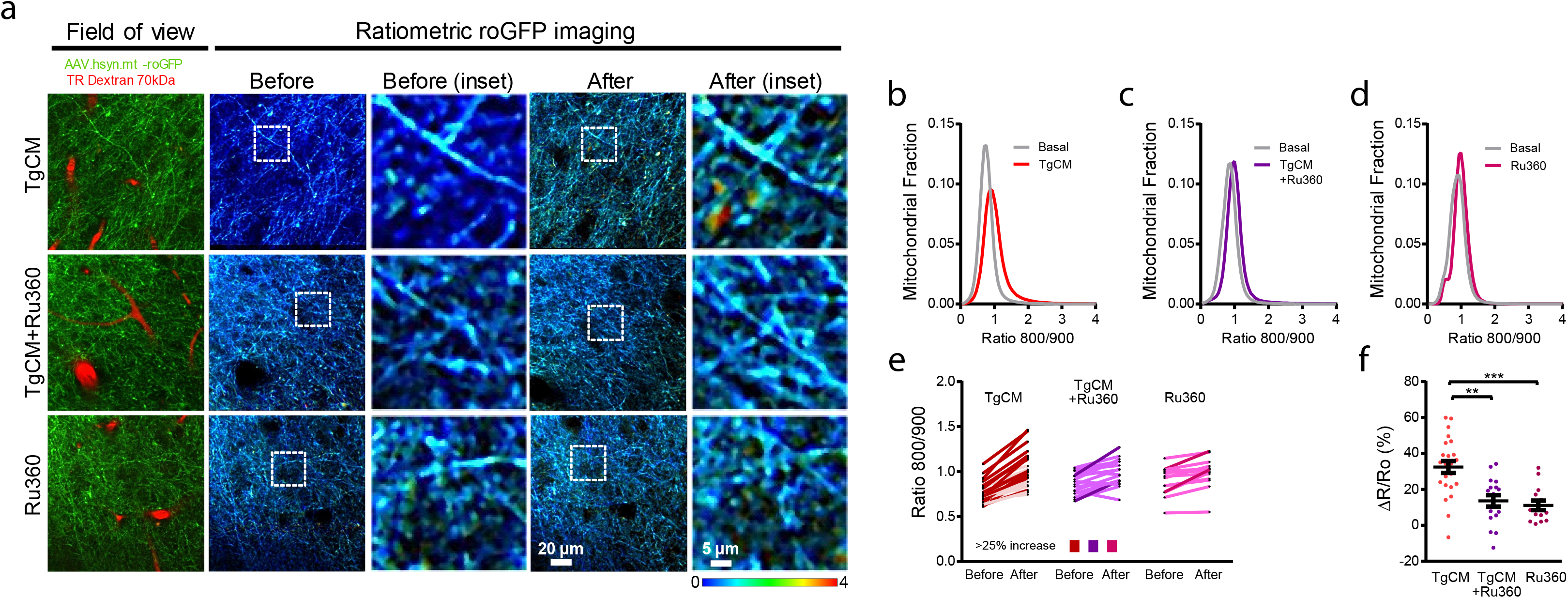
Inhibition of the MCU prevents neurons from mitochondrial oxidative stress. **a.** Representative pictures of the effects of the inhibition of the MCU on the mitochondrial oxidative stress induced by TgCM in neurons. Scale bar represents 20 μm and 5 μm in the insets. After exposure to Ru360, TgCM not was able to increase oxidative stress in mitochondria. **b - d.** Histograms of mitochondrial oxidative stress frequency distribution (Ratio 800/900) before and after application of TgCM with or without Ru360, and Ru360 alone. **e.** Averaged 800/900 ratios before and after TgCM treatment with or without Ru360, and Ru360 alone for each z-stack acquired. Darker traces represent z-stacks showing an increase ≥ 25% in 800/900 ratio after topical application of TgCM or Ru360. **f.** Scatter dot plot shows the relative change (ΔR/Ro) in ratio 800/900 for each condition. Error bars represent mean ± SEM (TgCM: 32.50 ± 3.328%, 24 z-stack; TgCM + Ru360: 13.64 ± 3.187%, 17 z-stacks; Ru360: 11.14 ± 2.660%, 14 z-stacks from 4, 4 and 3 mice respectively, **p<0.01, ***p<0.001).

### 5. Aβ-associated mitochondrial oxidative stress is reduced with the mitochondrial-targeted antioxidant SS31 in AD transgenic mouse neurons

To further evaluate the value of mitochondrial oxidative stress as a therapeutic target, we next evaluated the efficacy of SS31, a mitochondria-targeted antioxidant (31, 32), to reduce the mitochondrial oxidative stress levels in APP/PS1 Tg mice. Elamipretide (SS31, D-Arg-Dmt-Lys-Phe-NH2) peptide (31, 32), is a mitochondrial targeted antioxidant reported to eliminate ROS and increase ATP production in mitochondria, thus maintaining the mitochondrial membrane potential, and preventing the opening of the mitochondrial permeability transition pore, mitochondrial swelling and cytochrome c release (33). SS31 is currently being tested at the preclinical state as a promising drug against neurodegenerative diseases, inflammatory diseases, and ischemia-reperfusion injury (34–40).

APP/PS1 Tg mice and non-Tg controls were intraperitoneally injected for 8 weeks with 5 mg/kg SS31, starting at 8 mo of age. We used SS20, another SS tetra-peptide which lacks the free radical scavenging properties, as control (31). AAV.hSyn.mt-roGFP was intracortically injected, and mitochondrial redox levels in neurons were evaluated when mice were 10 mo old. Treatment with SS31 in APP/PS1 Tg mice led to a significant decrease in the relative mitochondrial oxidative stress (Ratio 800/900) relative to SS20-treated APP/PS1 Tg mice, and down to the levels of non-Tg mice treated with SS31 (Figure 5). Additionally, the statistically significant association between oxidative stress levels and the proximity to Aβ plaques was lost in SS31-treated APP/PS1 Tg mice (Figure 5e), as compared to the SS20-treated APP/PS1 Tg mice. These findings show that SS31 modifies mitochondrial redox state and ameliorates Aβ-related toxicity in neuronal mitochondria, holding promise for clinical trials in AD.

**Figure 5.**
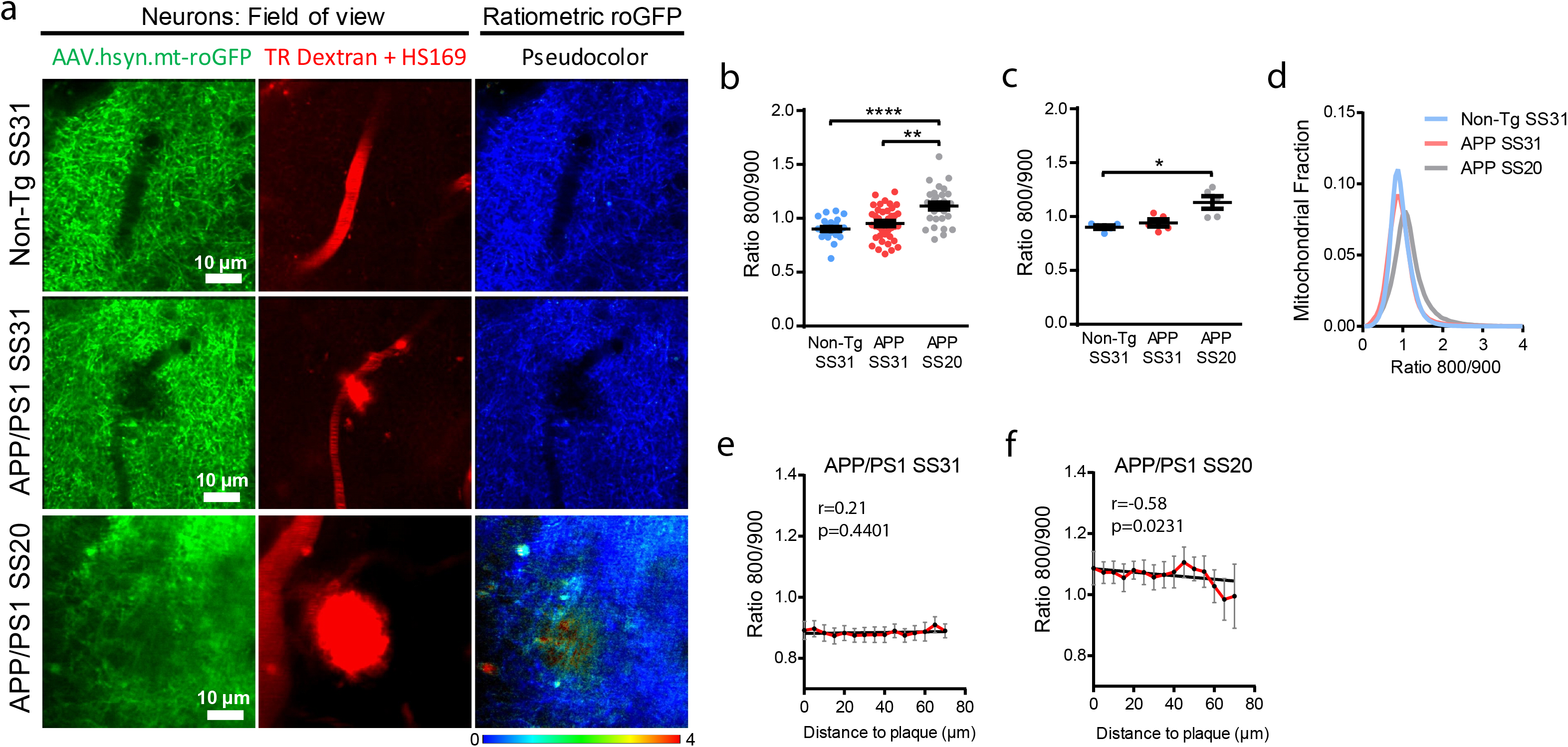
The antioxidant targeted to mitochondria SS31 reduces mitochondrial oxidative stress in AD transgenic mouse neurons. **a.** *In vivo* images of neurites and cell bodies expressing AAV.hSyn.mt-roGFP in mitochondria in non-Tg (top) and APP/PS1 Tg mice treated with either SS31 (middle) or SS20 (bottom). Field of view shows two-photon images of the expression of the AAV (green), blood vessels (Dextran, red) and amyloid plaques (HS169, red) in the cortex. Ratiometric roGFP imaging shows pseudocolor images according to the pseudocolor scale at the bottom. Scale bar represents 10 μm. **b, c.** Mitochondrial oxidative stress (Ratio 800/900) comparison between non-Tg and APP/PS1 Tg mice injected with either SS31 or SS20 in neuronal mitochondria (**b**. Average per field of view, non-Tg SS31: 0.94 ± 0.023, n = 32 z-stacks; APP/PS1 SS31: 0.95 ± 0.024, n = 40 z-stacks; APP/PS1 SS20: 1.11 ± 0.032, n = 30 z-stacks from 6, 6 and 5 mice respectively, ***p = 0.0001, **p<0.01. **c**. Average per mouse, non-Tg SS31: 0.90 ± 0.016, n = 6 mice; APP/PS1 SS31: 0.94 ± 0.032, n = 6 mice; APP/PS1 SS20: 1.13 ± 0.058, n = 5, *p <0.05). Error bars represent mean ± SEM. **d**. Histogram of mitochondrial oxidative stress frequency distribution (Ratio 800/900) of the three conditions. **e, f**. For APP/PS1 Tg mice injected with SS31, the probability of finding mitochondrial oxidative stress in neurons was similar at any distance within 70 µm from the edge of a plaque, unlike SS20 (mean ± SEM; APP/PS1 SS31 *r* (Pearson’s coefficient) 0.21, R square 0.046, n.s., n = 56 plaques from 6 Tg mice; APP/PS1 SS20 *r* −0.58, R square 0.338, *p<0.05, n = 19 plaques analysed from 4 Tg mice).

To determine whether SS31 reduces Aβ plaque burden, we quantified the extent of Aβ deposition in the cortex of SS31- and SS20-treated APP/PS1 Tg mice *ex vivo*. Conventional anti-Aβ immunolabeling detecting all Aβ deposits (diffuse and dense-core plaques) and ThioS staining (detecting the dense-core of plaques) revealed that neither the Aβ area fraction nor the density of Aβ deposits was significantly different between SS31- and SS20-treated APP/PS1 Tg mice (Supplemental Figure 5a, b). Mitochondria are highly impaired in the dystrophic neurites that surround amyloid plaques (41, 42). Dystrophic neurites present high levels of oxidation, especially surrounding Aβ plaques ((43)). To further evaluate the protective effects of SS31 in the Aβ plaque microenvironment, we analyzed the density of neuritic dystrophies per plaque (21, 44, 45). Numerous abnormal neurites were generally associated with amyloid plaques in SS20-treated APP/PS1 Tg mice, but the total amount of dystrophies as well as the percentage of plaques showing dystrophies were reduced in Tg mice treated with SS31 (Supplemental Figure 5c-e). Taken together, these results show the dramatic impact of SS31 on an array of Aβ-associated neurotoxic events in the vicinity of Aβ plaques.

### 6. Human brain RNA-seq shows a downregulation of neuronal mitochondrial antioxidant capacity in AD

Finally, we evaluated the mitochondrial antioxidant defense capacity of neuronal mitochondria in the AD vs. the normal aging brain by interrogating publicly available human single-nuclei RNA-seq dataset (46). Specifically, we compared the expression of 31 genes encoding antioxidant mitochondrial proteins in various cell types across AD (Braak NFT stages V/VI) and control (Braak stages 0/I/II) individuals. We observed that the expression of mitochondrial antioxidant defense genes was downregulated in neurons (both excitatory and inhibitory), with no changes observed in astrocytes and other cell types (oligodendrocytes and endothelial cells) (Figure 6 and Supplemental Table 1). These results suggest that the expression of most human genes involved in the mitochondrial antioxidant defense of neurons is altered in AD patients at the transcriptional level.

**Figure 6.**
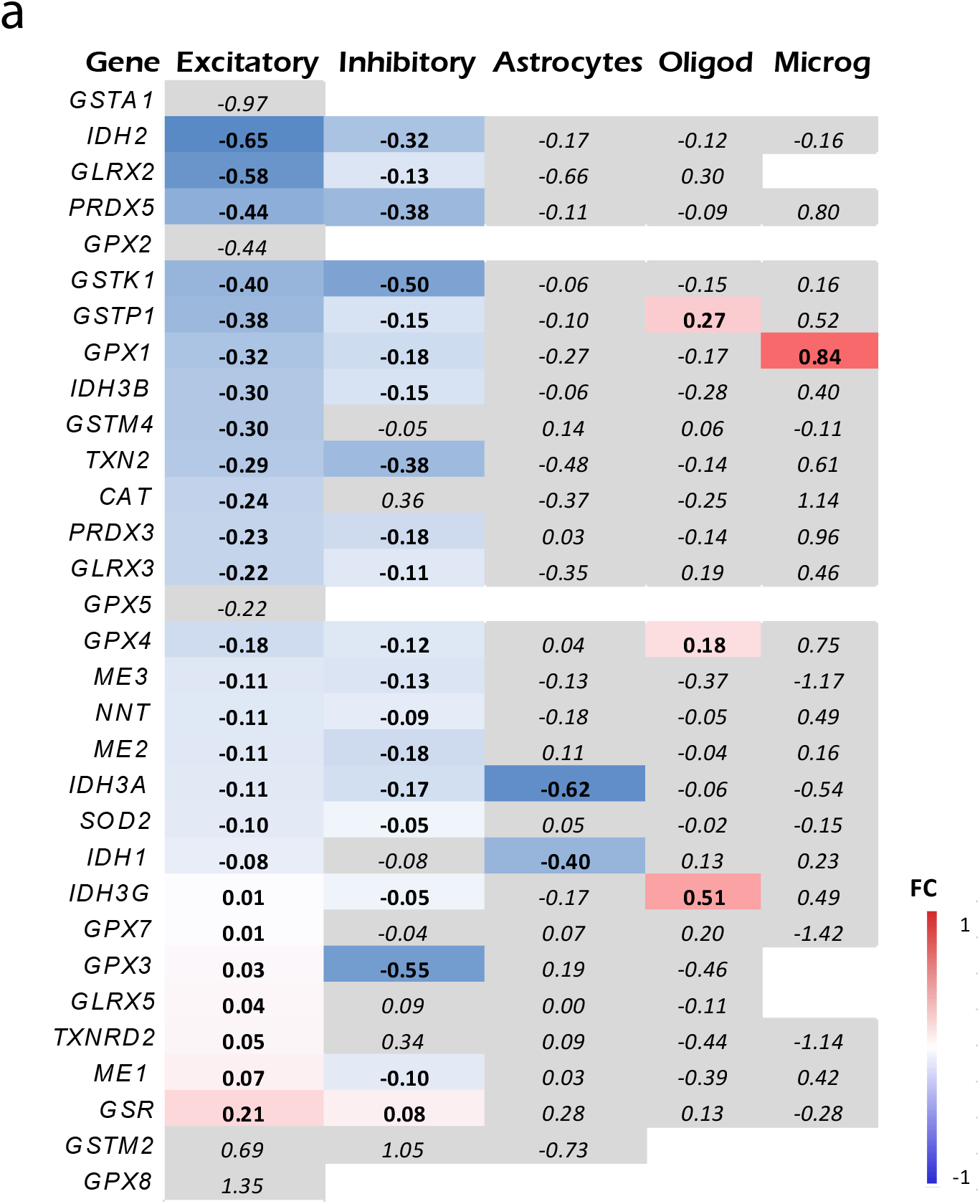
Single cell RNA-seq shows a decrease in the mitochondrial antioxidant defense in neurons. **a.** Heat map representing fold change (FC) of the expression of genes encoding proteins involved in mitochondria antioxidant defense between AD (Braak NFT stages V/VI) and control individuals (Braak NFT stages 0/I/II) (from (46)). The color of each box indicates the direction and magnitude of gene expression fold change. Grey color means statistically non-significant (p>0.05) differences. Note that the expression of mitochondrial antioxidant defense genes is overall downregulated in both excitatory and inhibitory neurons, whereas no change is observed in astrocytes and other cell types. *CAT* (catalase), *GLRX* (glutaredoxin), *GPX* (glutathione peroxidase), *GSR* (glutathione reductase), *GST* (glutathione transferase), *IDH* (isocitrate dehydrogenase), *PRDX* (peroxiredoxins), *ME* (malic enzyme), *NNT* (nicotinamide nucleotide transhydrogenase), *SOD2* (manganese superoxide dismutase), *TXN2* (thioredoxin), *TXNRD* (thioredoxin reductase).

## Discussion

Previous *in vitro* and *in vivo* studies suggest that mitochondrial oxidative stress is related to AD pathology (47–51). However, the sequence of pathological events was not well understood in the *in vivo* living brain, as well as whether the use of mitochondrial antioxidant targeted compounds could prevent these effects. Here, we address these questions by using multiphoton microscopy in combination with cell type-specific redox analysis in a mouse model of cerebral β amyloidosis. The main findings of our *in vivo* study are as follows: (1) Mitochondrial oxidative stress is exacerbated in neurons in surrounding Aβ plaques; (2) soluble Aβ oligomers contribute to the increased mitochondrial oxidative stress levels; (3) Aβ-driven increased mitochondrial redox levels can be corrected by precluding mitochondrial Ca^2+^ influx into mitochondria via MCU inhibition; (4) the mitochondrial targeted antioxidant SS31 prevents Aβ-induced increased oxidative stress levels in neuronal mitochondria.

We found increased mitochondrial redox levels in the APP/PS1 Tg mouse only after Aβ plaque deposition, and particularly close to plaques. Our group has previously shown that oxidative stress in the neuronal cytosol also occurs after plaque deposition in the APP/PS1 Tg mouse (43), and our present findings suggest that this feature could be related secondary to mitochondrial damage. Neurons are highly compartmentalized cells, and every compartment can have unique mitochondrial requirements (52). We observed this phenomenon both in neurites and in somas, although the stress levels were consistently higher in the neurites. Dendrites are usually more susceptible to oxidative insult (53), and damaged mitochondria are known to have reduced motility, minimizing the spread of oxidation throughout the neuron and especially the soma, possibly explaining these differences between neurites and somas. Alternatively, this finding could be related to an increased bioenergetic demand associated with buffering Ca^2+^ near a synapse (54).

In line with current results, our group has previously shown structurally and functionally damaged mitochondria around Aβ plaques (55). These mitochondria are prone to produce more ROS and less ATP, likely contributing to neurodegenerative phenotypes observed in these mice such as dystrophic neurites and synaptic loss. Herein, a major focus of our study was to analyze the impact of acute oxidative insult induced by soluble Aβo within the neuronal mitochondria. We found that mitochondrial redox state levels were increased after application of soluble Aβo (TgCM) to the naïve Wt mouse brain, (vs. WtCM or Aβ-immunodepleted TgCM), implying that Aβo are also involved in the neuronal mitochondrial oxidative stress observed in AD. In line with these results, the concentration of soluble oligomeric Aβ increases with age in the APP/PS1 Tg mouse model (16, 56, 57), which could contribute to the increased redox level observed in the old Tg mice.

Increased neuronal activity enhances mitochondrial superoxide production via excessive cytosolic Ca^2+^ load (58–60), which could in turn lead to mitochondrial Ca^2+^ overload, since mitochondria buffer cytosolic Ca^2+^ leading to neurotoxicity. We found that inhibition of MCU, the main route for Ca^2+^ influx into the mitochondria, via blocking its pore with the specific drug Ru360, attenuated the Aβ-driven mitochondrial oxidative stress, which places mitochondrial Ca^2+^ dyshomeostasis upstream of oxidative stress, and deciphers the most likely sequence of events *in vivo*. While dysregulation in mitochondrial Ca^2+^ signaling and increased ROS production have been observed separately in various *in vitro* models and a *C. elegans* model of AD (61), here we have demonstrated a link between both deleterious events *in vivo* in a mouse model of AD. Future work will determine whether Aβ drives a vicious cycle in which mitochondrial Ca^2+^ overload leads to mitochondrial oxidative stress and this exacerbates mitochondrial Ca^2+^ overload, since a better understanding of the links between mitochondrial Ca^2+^, MCU, and oxidative stress could propel the development of mitoprotective therapies against AD and perhaps other neurodegenerative diseases.

Neurons need to maintain mitochondrial function even after a local insult, and to accomplish this, the cells use mitochondrial antioxidants which protect against the ensuing toxicity. Severely damaged mitochondria decrease their antioxidant defense, thus increasing ROS production. In line with this concept, we observed a downregulation of the expression of the mitochondrial antioxidant defense genes in neurons from AD brains by interrogating a public single-nuclei RNA-seq dataset. Additionally, the mitochondrial antioxidant defense could be impaired in the setting of mitochondrial Ca^2+^ overload due to the direct inhibition of the antioxidant enzymes due to the high Ca^2+^ levels (62).

Using the hSyn.mt-roGFP sensor and multiphoton microscopy, we also found relative reduced mitochondrial oxidative stress in neurons in the SS31-treated APP/PS1 Tg mice, compared to SS20-treated ones. Therapeutic clinical trials with natural antioxidants have generally shown mixed results. Some prevention trials have shown that treatment of elderly people with vitamins C and E reduces the risk of AD (63–65), whereas others did not (66–69). It has been proposed that these hydrosoluble vitamins are not able to cross the blood brain barrier and reach brain cells. SS31 is a cell permeable small peptide that can easily penetrate in neurons and concentrate in the mitochondrial matrix (70). SS31 has been shown to protect against Ca^2+^-induced mitochondrial depolarization and swelling, and several mitochondrial insults, including Aβ toxicity, in treated N2a cells and primary neurons from Tg mice (71). While another study reported that SS31 reduces soluble Aβ levels and Aβ deposits in APP Tg mice, leading to an improvement in mitochondrial function (72), we did not detect significant changes in Aβ burden, whether measured by area fraction (% immunoreactive area) or density (number/mm^2^) of total or dense-core (ThioS-positive) plaques. Instead, the number of dystrophic neurites per plaque was significantly reduced upon SS31 treatment, suggesting that mitochondrial oxidative stress contributes to this neurodegenerative feature of Aβ plaques (73). This finding is particularly intriguing because Aβ plaque-associated dystrophic neurites are known to accumulate damaged mitochondria (42, 74).

## Conclusion

Taken together, our findings expand the current knowledge about mitochondrial redox homeostasis in neurons and its impairment in AD. By combining a mitochondrial redox reporter and multiphoton microscopy, we have shown that neurons exhibit exacerbated mitochondrial oxidative stress levels in the presence of Aβ aggregates *in vivo*. Even though we are aware that a change in the mt-roGFP ratio does not mandate ROS generation in mitochondria (since ROS could be also generated in the cytosol and diffuse later into mitochondria), this sensor provides a measurement of the relative mitochondrial oxidative stress regardless of its source. We have also provided evidence that mitochondria-targeted antioxidants can ameliorate mitochondrial Aβ-induced oxidative stress in neurons, and might represent a novel strategy with potential therapeutic and/or preventative value in AD.

## Abbreviations

AAV: adeno-associated virus
AD: Alzheimer’ disease
Aβ: amyloid beta
CM: conditioned media
DTDP: dithiodipyridine;
DTT: dithiothreitol
MCU: mitochondrial calcium uniporter
mt-roGFP: matrix-roGFP
ROS: reactive oxygen species
SS31: Elamipretide
Tg: transgenic
Wt: wild-type

## Acknowledgements

Authors acknowledge the Schepens Eye Research Institute Gene Transfer Vector Core for the AAV.hSyn.mt-roGEP vector production.

## Author contributions

MCR designed experiments, collected and analysed data, and wrote the original draft. EKK, ACS, and MVSM helped with immunohistochemistry and mouse brain injections. SSH wrote ImageJ and Matlab macros. SD analysed single RNA-seq data. ZF performed cloning design. HS and KPRN provided LCOs. ASP provided expertise and feedback and edited the manuscript. BJB conceptualized the research, designed experiments, discussed data, edited the manuscript and secured funding. All authors read and approved the final manuscript.

## Data and materials availability

The authors declare that all data supporting the findings of this study are available within the article and its Supplementary Information files or from the corresponding author upon reasonable request.

## Supplementary materials

**Fig. S1. Validation of AAV.hSyn.mt-roGFP *in vitro*.**

**Fig. S2. Validation of pAAV.hSyn.mt-roGFP *ex vivo*.**

**Fig. S3. Neurites show increased mitochondrial oxidative stress levels.**

**Fig. S4. Mitochondrial oxidative stress is not elevated in AD transgenic mouse neurons before Aβ plaque deposition.**

**Fig. S5. SS31 reduces Aβ-associated dystrophic neurite number but not Aβ burden in APP/PS1 Tg mouse.**

**Supplemental Table 1. Analysis of neuronal mitochondrial antioxidant capacity in AD vs. normal aging brain.**

