## Supplemental information for "Real-time imaging of mitochondrial redox reveals increased mitochondrial oxidative stress associated with amyloid β aggregates *in vivo* in a mouse model of Alzheimer’s disease"

**This PDF file includes:**

Supplementary text

Figures S1 to S5

Tables S1

**Fig. S1. Validation of AAV.hSyn.mt-roGFP *in vitro.* a.** Mitochondrial co-transfection verified proper targeting of mt-roGFP to mitochondria. N2a cells (top) and primary cortical neurons (bottom) were co-transfected with mt-roGFP (green) and mRuby-Mito-7 (red) and subjected to confocal microscopy imaging. Scale bar represents 10 μm. **b.** Double immunolabelling of mt-roGFP (green) and mRuby-ER5 (red, targeting endoplasmic reticulum, ER) in N2a cells shows lack of colocalization and supports the mitochondrial localization of mt-roGFP. Scale bar represents 10 μm. **c.** *In vitro* imaging of cellular oxidative stress with mt-roGFP. Primary cortical neurons were exposed to either the oxidant DTDP or the reducing agent DTT. Images at 800 nm (red), 900 nm (green) and merged are shown. **d.** The relative changes in ratio 800/900 were represented by histograms of ratio 800/900 frequency distribution in control conditions (grey) and 20 min after exposure to DTT 1 mM (blue) and DTDP 100 μM (red) (Control, n = 143 cells; DTT 1 mM, n = 125; DTDP 100 μM, n = 109 cells).

**Fig. S2. Validation of pAAV.hSyn.mt-roGFP *ex vivo.*** AAV.hSyn.mt-roGFP targets neuronal mitochondria *in vivo.* **a**. Colocalization of AAV.hSyn.mt-roGFP (green), NeuN (red) and GS (glutamine synthetase, magenta) in the mouse cortex shown by immunohistochemistry. Note that AAV.hSyn.mt-roGFP only colocalizes with the neuronal marker NeuN. **b**. Colocalization of AAV.hSyn.mt-roGFP (green), HSP60 (mitochondrial marker, red) and NeuN (magenta) in the cortex shown by immunohistochemistry. **c.** Inset. Colocalization of AAV.hSyn.mt-roGFP (green) and HSP60 (red) in cortex shown by immunohistochemistry (top). Scale bar 5 μm. Graph shows intensity profile of the ROI across the cell. Green line represents the fluorescence intensity of AAV.hSyn.mt-roGFP and red line represents the fluorescence intensity of HSP60.

**Fig. S3. Neurites show increased mitochondrial oxidative stress levels.** Comparison of mitochondrial oxidative stress (Ratio 800/900) in the different cell compartments (somas and neurites) in 10-month-old (old) and 3-month-old (young) non-Tg and APP/PS1 Tg mice. Neurites showed significantly higher oxidative stress levels in mitochondria at both ages and conditions. Error bars represent mean ± SEM. (**a**. Old non-Tg: 0.87 ± 0.024 for somas and 0.99 ± 0.039 for neurites, n = 13 z-stacks from 4 mice, ***p = 0.0003. **b**. Old APP/PS1: 1.08 ± 0.065 for somas and 1.35 ± 0.084 for neurites, n = 12 z-stacks from 6 mice, ****p < 0.0001. **c**. Young non-Tg: 0.76 ± 0.028 for somas and 0.86 ± 0.045 for neurites, n = 6 z-stacks from 2 mice, *p = 0.031. **d**. Young APP/PS1: 0.75 ± 0.026 for somas and 0.92 ± 0.030 for neurites, n = 10 z-stacks from 3 mice, ***p = 0.0003).

**Fig. S4. Mitochondrial oxidative stress is not elevated in AD transgenic mouse neurons before Aβ plaque deposition. a.** *In vivo* images of neurites and cell bodies expressing pAAV.hSyn.mt-roGFP in mitochondria in non-Tg (top) and APP/PS1 Tg mice (bottom) in young mice. Scale bar represents 10 μm. **b, c.** Scatter dot plot represents mitochondrial oxidative stress (Ratio 800/900) in non-Tg and APP/PS1 Tg mice at 3 months of age, before plaque deposition, in mitochondria in neurons (**b**, average per field of view, non-Tg: 0.83 ± 0.024, n = 18 z-stacks; APP/PS1: 0.87 ± 0.024, n = 42 z-stacks from 3 and 6 mice respectively; **c**. average per mouse, non-Tg: 0.82 ± 0.039, n = 3 mice; APP/PS1: 0.87 ± 0.034, n = 6 mice). Error bars represent mean ± SEM. **c**. Histogram of mitochondrial oxidative stress frequency distribution (indicated by Ratio 800/900) in the young non-Tg and APP/PS1 Tg mice. **d, e**. Comparison of mitochondrial oxidative stress (Ratio 800/900) in somas and neurites in 3-month-old non-Tg and APP/PS1 Tg mice. No differences were found. Error bars represent mean ± SEM. (**d**. somas: 0.77 ± 0.028, n = 6 z-stacks from 2 non-Tg mice, and 0.75 ± 0.026, n = 10 z-stacks from 3 APP/PS1 Tg mice. **e**. neurites: 0.87 ± 0.045, n = 6 z-stacks from 2 Non-Tg mice, and 0.92 ± 0.030, n = 10 z-stacks from 3 APP/PS1 Tg mice).

**Fig. S5. SS31 reduces Aβ-associated dystrophic neurite number but not amyloid burden in the AD transgenic mouse. a.** Representative images of the global amount of amyloid in the cortex of SS31 and SS20 treated APP/PS1 mice at 10 mo of age after Aβ immunostaining. **b.** Scatter dot plots represent the quantification of amyloid load in the cortex after anti-Aβ immunostaining or ThioS labeling. The amount of dense-core plaques detected by ThioS (top) and the overall load of Aβ (bottom) was comparable among SS31 and SS20 APP/PS1 treated mice. n = 7 mice per condition. Histograms represent the dense core plaque (top) and diffuse amyloid deposit (bottom) size in both conditions. **c.** Representative images of neuritic dystrophies (arrow heads, neurofilaments in green) around amyloid plaques (blue) in APP/PS1 mouse after either SS31 or SS20 treatment. Scale bar 20 μm. **d.** Scatter dot plot represents the quantification of the number of dystrophic neurites observed per plaque, n = 362 plaques from 4 SS31 APP/PS1 treated mice and n = 295 plaques from 4 SS31 APP/PS1 treated mice, **p < 0.05. **e.** Scatter dot plot represents the percentage of plaques showing dystrophic neurites, n = 4 – 5 areas per 4 mouse per condition, *p = 0.022.

**Supplemental Table 1. Analysis of neuronal mitochondrial antioxidant capacity in AD vs. normal aging brain.** The expression levels for genes encoding antioxidant enzymes (CAT, GLRX, GPX, GSR, GST, IDH, PRDX, ME, NNT, SOD2, TXN2, and TXNRD) were compared between control (B1, Braak NFT stages 0/I/II) and AD (B3, Braak NFT stages V/VI) individuals of a publicly available human single-nuclei RNA-seq [44]. The average expression level for each gene and group are shown, together with the log fold change and the adjusted p-value of the individual gene models, and the z-scores and the p-values of the mixed models.
