## Supplemental figures for "Real-time imaging of mitochondrial redox reveals increased mitochondrial oxidative stress associated with amyloid β aggregates *in vivo* in a mouse model of Alzheimer’s disease"

Supplementary Figure 1.

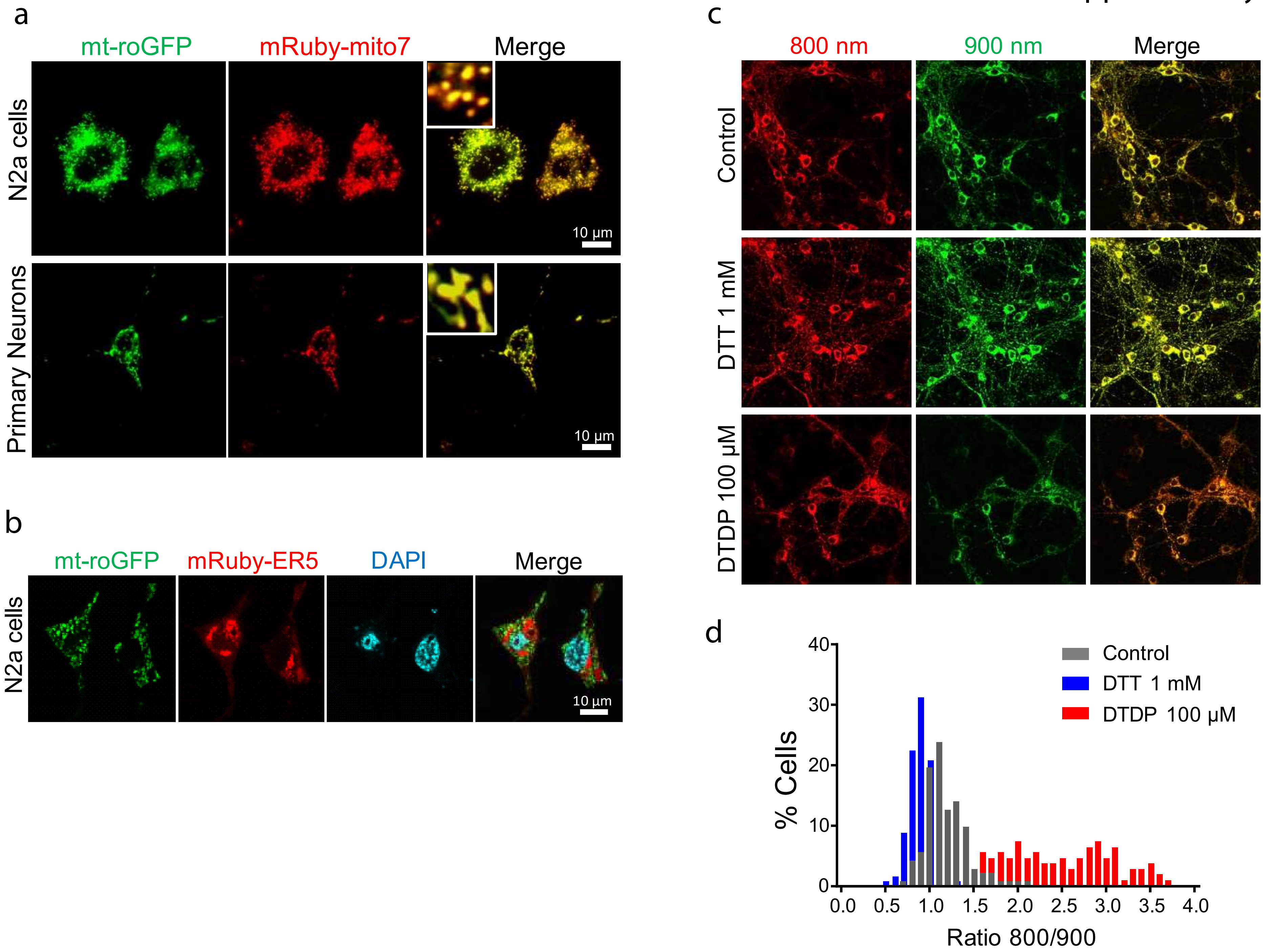

Supplementary Figure 2.

a

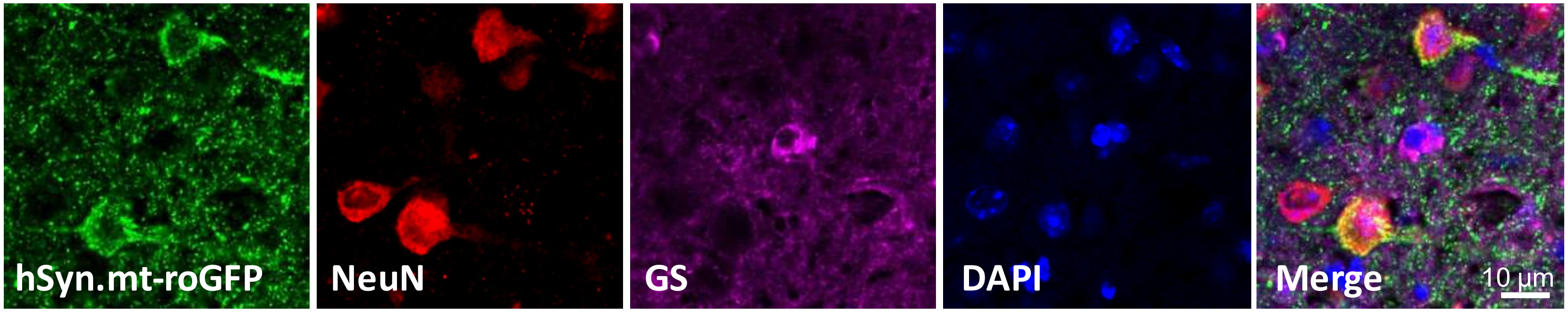


b

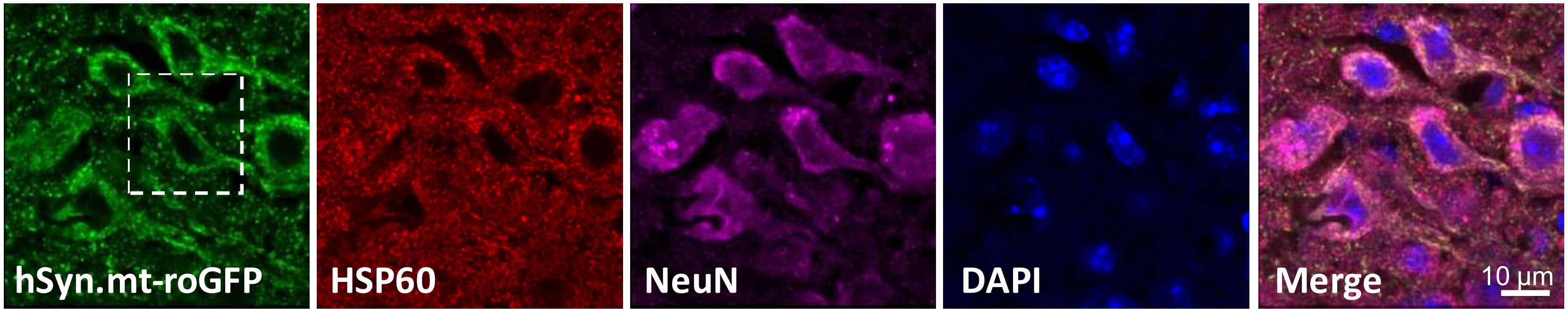


c

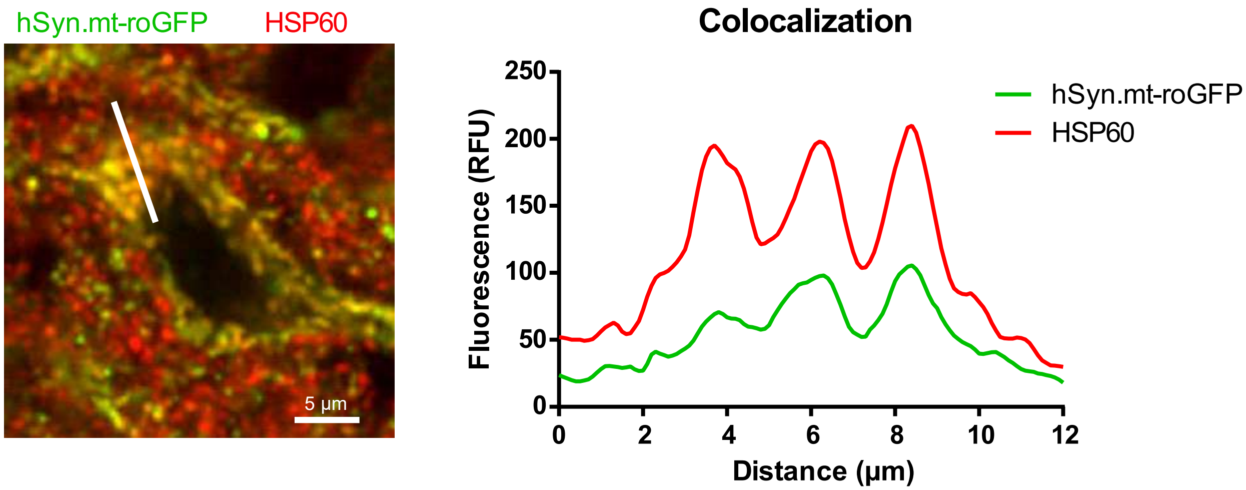

Supplementary Figure 3.

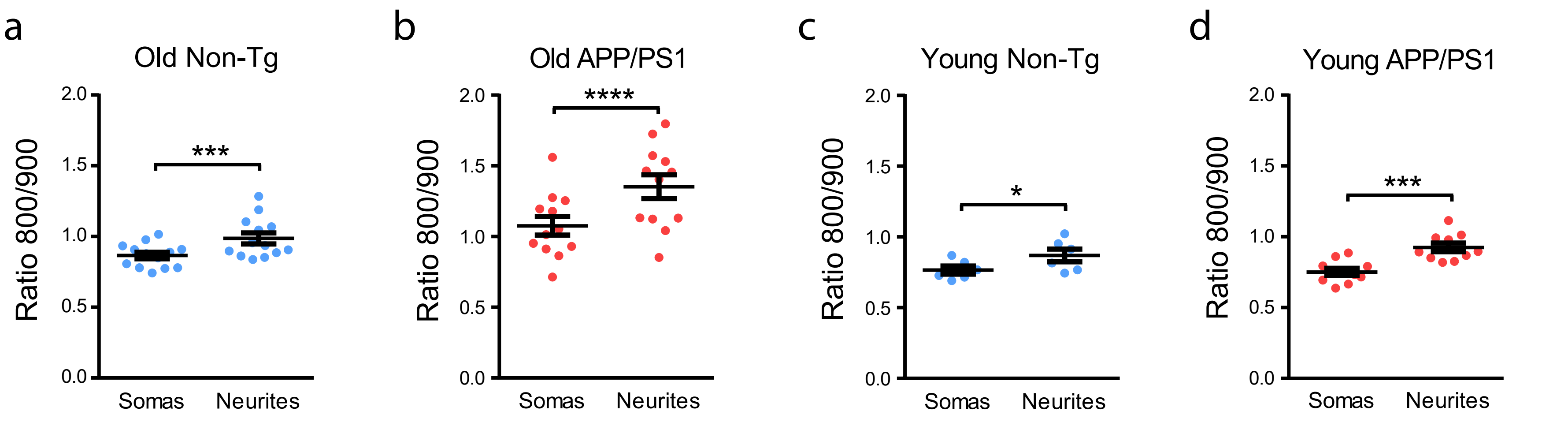

Supplementary Figure 4.

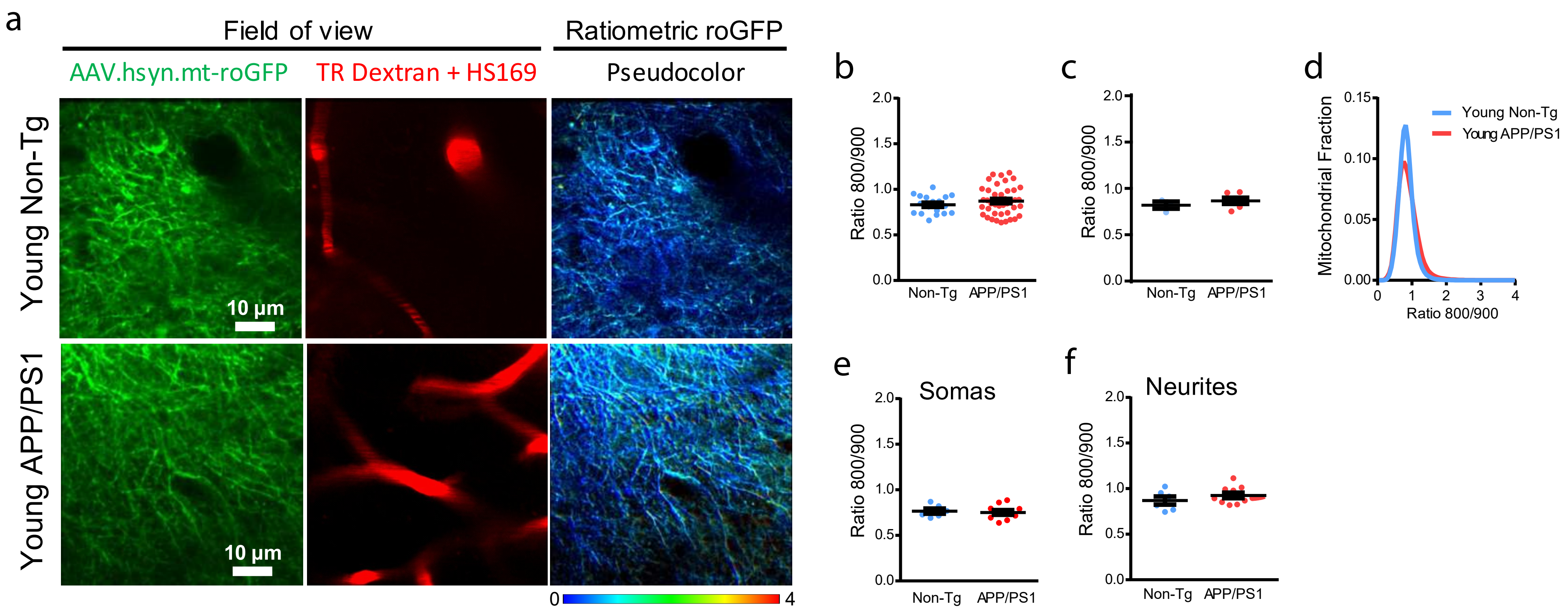

Supplementary Figure 5.

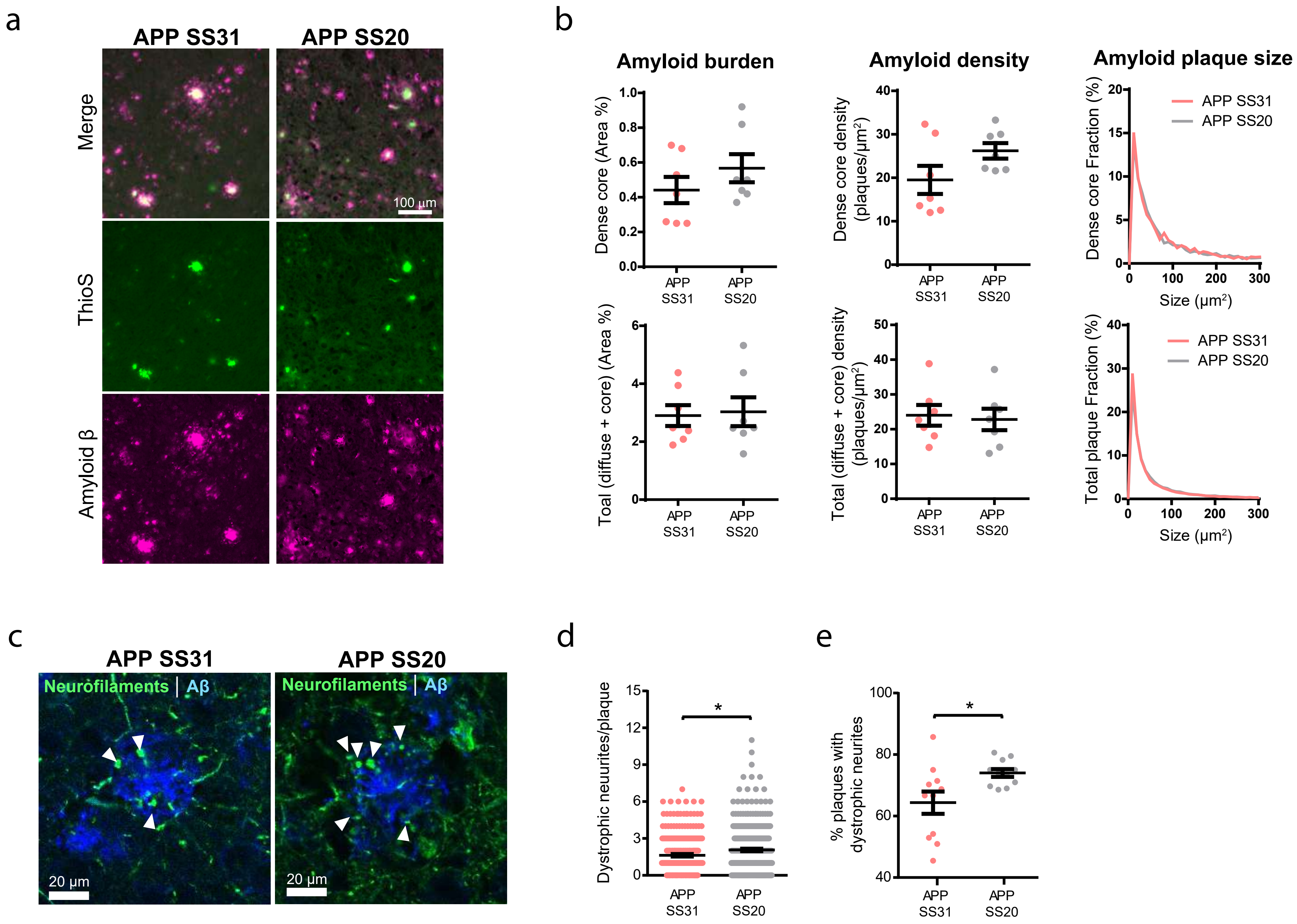

| Cell type | Gene | CTRL mean | AD mean | log FG | Adj. p-value | z-score | p-value |
|---|---|---|---|---|---|---|---|
| | IDH2 | 0.18796272 | 0.1197171 | -0.65082 | 1.33E-133 | -10.9863 | 4.45E-28 |
| | PRDX5 | 0.27819673 | 0.204403 | -0.44469 | 1.25E-106 | -4.58978 | 4.44E-06 |
| | GPX1 | 0.3278097 | 0.2628929 | -0.31838 | 1.56E-84 | -2.5134 | 0.011957249 |
| | ME3 | 0.43422281 | 0.4009608 | -0.11497 | 2.54E-79 | -3.47341 | 0.000513891 |
| | GPX4 | 0.6085328 | 0.5371614 | -0.17998 | 8.80E-75 | -0.77569 | 0.437931461 |
| | NNT | 0.46874834 | 0.433145 | -0.11396 | 1.92E-65 | -1.36825 | 0.171234334 |
| | GSTP1 | 0.16003103 | 0.1225748 | -0.38469 | 7.86E-63 | -2.4275 | 0.015203459 |
| | ME2 | 0.22026604 | 0.2041281 | -0.10977 | 2.43E-60 | -3.07736 | 0.002088423 |
| | IDH3A | 0.21737885 | 0.2019876 | -0.10594 | 1.65E-56 | -2.17463 | 0.029657966 |
| | GSTK1 | 0.12504372 | 0.094587 | -0.40272 | 4.85E-56 | -3.78133 | 0.000155993 |
| | SOD2 | 0.49597773 | 0.4634756 | -0.09778 | 6.64E-52 | -0.81511 | 0.415012183 |
| | TXN2 | 0.12350612 | 0.1011846 | -0.28759 | 8.45E-48 | -2.87726 | 0.004011492 |
| | GLRX3 | 0.14323475 | 0.122712 | -0.22311 | 2.48E-46 | -3.07693 | 0.00209141 |
| | PRDX3 | 0.18441932 | 0.1577318 | -0.22552 | 8.03E-45 | -3.1825 | 0.001460098 |
| | IDH3B | 0.11448413 | 0.0929703 | -0.30031 | 2.59E-42 | -1.58472 | 0.113029374 |
| Excitatory | GLRX2 | 0.08399903 | 0.0560591 | -0.58342 | 4.92E-39 | -3.66377 | 0.000248529 |
| | ME1 | 0.40133441 | 0.421476 | 0.07065 | 9.49E-31 | 2.061165 | 0.039287263 |
| | GSTM4 | 0.05885529 | 0.0478255 | -0.29939 | 3.67E-27 | -1.52494 | 0.127273589 |
| | GPX3 | 0.13046523 | 0.133474 | 0.03289 | 9.14E-27 | -1.35877 | 0.174218862 |
| | CAT | 0.04904186 | 0.0416225 | -0.23665 | 6.34E-26 | -3.25874 | 0.001119088 |
| | IDH3G | 0.13565684 | 0.1365158 | 0.00911 | 3.51E-23 | 1.22953 | 0.218873103 |
| | IDH1 | 0.08891133 | 0.0841164 | -0.07998 | 1.08E-17 | 1.693496 | 0.090361039 |
| | TXNRD2 | 0.08043839 | 0.0830085 | 0.04538 | 1.31E-16 | 0.358492 | 0.719975127 |
| | GLRX5 | 0.07313464 | 0.0749886 | 0.03612 | 5.01E-14 | -0.03955 | 0.968449793 |
| | GSR | 0.11559692 | 0.1333014 | 0.20559 | 1.16E-08 | 3.475353 | 0.000510182 |
| | GPX7 | 0.00795432 | 0.0080109 | 0.01023 | 0.042385977 | 1.311759 | 0.189601582 |
| | GSTA1 | 0.00104158 | 0.0005335 | -0.96511 | 0.092555287 | | |
| | GSTM2 | 0.00400242 | 0.0064438 | 0.68704 | 0.228458887 | | |
| | GPX2 | 0.00068763 | 0.0005057 | -0.44346 | 0.482749683 | | |
| | GPX5 | 0.00017517 | 0.0001506 | -0.21761 | 0.5417367 | | |
| | GPX8 | 0.00058334 | 0.0014902 | 1.35311 | 0.79809893 | | |
| | PRDX5 | 0.13734908 | 0.1053599 | -0.38252 | 5.92E-12 | -2.80445 | 0.005040225 |
| | ME3 | 0.32228437 | 0.2941451 | -0.13181 | 3.38E-10 | -0.75977 | 0.44739339 |
| | ME2 | 0.17949136 | 0.1579758 | -0.18421 | 2.10E-09 | -3.53084 | 0.000414247 |
| | GPX4 | 0.30882569 | 0.2842658 | -0.11955 | 3.17E-08 | 1.254055 | 0.209822085 |
| | GPX1 | 0.1665413 | 0.1469012 | -0.18103 | 2.15E-07 | -0.64837 | 0.516744641 |
| | IDH2 | 0.08452088 | 0.0675409 | -0.32355 | 2.44E-07 | -2.31899 | 0.020395839 |
| | ME1 | 0.18729903 | 0.1748807 | -0.09897 | 1.18E-06 | -0.53023 | 0.595955205 |
| | IDH3A | 0.13516119 | 0.1205188 | -0.16542 | 1.23E-06 | -1.58808 | 0.112268432 |
| | TXN2 | 0.07082787 | 0.0545646 | -0.37635 | 7.98E-06 | -1.1482 | 0.250884279 |
| | NNT | 0.21685105 | 0.2036071 | -0.09092 | 8.59E-06 | -1.34212 | 0.17955607 |
| | GSTP1 | 0.08835062 | 0.0798887 | -0.14525 | 0.000135905 | -1.73755 | 0.082289348 |
| | SOD2 | 0.20942942 | 0.2029384 | -0.04542 | 0.00039807 | -0.27649 | 0.782172837 |
| | GSTK1 | 0.08308578 | 0.0585545 | -0.50482 | 0.000581944 | -1.49802 | 0.134128064 |
| Inhibitory | GLRX2 | 0.03123418 | 0.0285408 | -0.1301 | 0.001961895 | -1.99102 | 0.046479158 |

| | | | | | | | |
|---|---|---|---|---|---|---|---|
| | PRDX3 | 0.09070094 | 0.0799008 | -0.18291 | 0.005236721 | -0.00184 | 0.998529032 |
| | IDH3G | 0.06376152 | 0.0615853 | -0.0501 | 0.005323398 | 0.370742 | 0.71082989 |
| | GLRX3 | 0.07647021 | 0.0708354 | -0.11043 | 0.009252403 | -0.71509 | 0.474553877 |
| | IDH3B | 0.05254184 | 0.0474246 | -0.14783 | 0.013509674 | 0.639959 | 0.52219906 |
| | GPX3 | 0.00576625 | 0.0039457 | -0.54735 | 0.023977277 | | |
| | GSR | 0.0977937 | 0.1031754 | 0.07729 | 0.030093706 | 1.380337 | 0.167483032 |
| | IDH1 | 0.06394805 | 0.0604522 | -0.0811 | 0.050863932 | 0.23363 | 0.815272155 |
| | CAT | 0.00461899 | 0.0059301 | 0.36049 | 0.35199375 | | |
| | GPX7 | 0.00855821 | 0.0083268 | -0.03955 | 0.361871949 | -0.48729 | 0.626049652 |
| | GSTM4 | 0.02951683 | 0.0284331 | -0.05397 | 0.463598174 | 0.6146 | 0.538819079 |
| | TXNRD2 | 0.04015504 | 0.0508591 | 0.34093 | 0.469745094 | 1.756618 | 0.078982971 |
| | GLRX5 | 0.02905416 | 0.03084 | 0.08606 | 0.48121826 | 0.46035 | 0.645265182 |
| | GSTM2 | 0.00461875 | 0.0095717 | 1.05127 | 0.539716633 | | |
| Astrocytes | IDH3A | 0.08392204 | 0.0545089 | -0.62256 | 0.001551929 | -3.29702 | 0.000977175 |
| | IDH1 | 0.13171162 | 0.0995762 | -0.40351 | 0.005924491 | -2.85945 | 0.004243736 |
| | CAT | 0.12526872 | 0.0972486 | -0.36528 | 0.085835655 | -2.32739 | 0.019944581 |
| | IDH2 | 0.27768046 | 0.2474353 | -0.16637 | 0.26595381 | -1.45367 | 0.146038852 |
| | GLRX3 | 0.06071736 | 0.0477207 | -0.34749 | 0.338147211 | -0.66807 | 0.504089083 |
| | TXN2 | 0.05051066 | 0.0361069 | -0.48431 | 0.344325586 | -0.84173 | 0.399938115 |
| | NNT | 0.14219869 | 0.1258381 | -0.17634 | 0.453013958 | -1.28954 | 0.197210975 |
| | GLRX2 | 0.01259626 | 0.0079494 | -0.66408 | 0.522432236 | -0.9931 | 0.320659117 |
| | PRDX3 | 0.02860987 | 0.0291534 | 0.02715 | 0.522432236 | -0.7694 | 0.441653452 |
| | GSTM2 | 0.02615143 | 0.0157208 | -0.73422 | 0.536901426 | -0.59881 | 0.549299657 |
| | IDH3B | 0.03330224 | 0.0319892 | -0.05804 | 0.599356128 | -0.04493 | 0.964161821 |
| | ME1 | 0.26234524 | 0.2686561 | 0.03429 | 0.62205411 | 1.132632 | 0.257368741 |
| | GPX1 | 0.03255749 | 0.0270267 | -0.26861 | 0.63958945 | -0.45992 | 0.645575118 |
| | ME2 | 0.11052652 | 0.1189926 | 0.10648 | 0.63958945 | 0.267556 | 0.789041367 |
| | ME3 | 0.07472281 | 0.0681268 | -0.13333 | 0.656218031 | -0.20121 | 0.840537686 |
| | GSTK1 | 0.06692031 | 0.0644037 | -0.0553 | 0.672679217 | 0.357342 | 0.720835701 |
| | GLRX5 | 0.01053364 | 0.0105307 | -0.0004 | 0.800546954 | 1.147767 | 0.25106481 |
| | TXNRD2 | 0.03424624 | 0.0364304 | 0.0892 | 0.810553048 | 0.989969 | 0.322189271 |
| | GPX4 | 0.09376679 | 0.096255 | 0.03778 | 0.873285048 | 0.293772 | 0.76893249 |
| | GSR | 0.02936822 | 0.0356514 | 0.2797 | 0.880819386 | 0.910486 | 0.362566449 |
| | PRDX5 | 0.07190518 | 0.066412 | -0.11465 | 0.907858173 | 1.287944 | 0.197765617 |
| | GSTP1 | 0.07882461 | 0.0733208 | -0.10442 | 0.919884111 | 0.217806 | 0.827580515 |
| | GPX3 | 0.00791244 | 0.0090243 | 0.18969 | 0.927001412 | | |
| | GSTM4 | 0.03402268 | 0.0375304 | 0.14156 | 0.929923259 | 0.958952 | 0.337582913 |
| | GPX7 | 0.01084284 | 0.0113625 | 0.06754 | 0.965931931 | 0.585103 | 0.558478493 |
| | IDH3G | 0.03122257 | 0.0277546 | -0.16986 | 0.970670192 | 0.382964 | 0.701746364 |
| | SOD2 | 0.15438949 | 0.1599526 | 0.05107 | 0.99359659 | 1.481806 | 0.138392017 |
| | GSTP1 | 0.07847924 | 0.0948515 | 0.27336 | 0.000216026 | 4.726363 | 2.29E-06 |
| | GPX4 | 0.06447126 | 0.0729387 | 0.17803 | 0.006160905 | 3.161555 | 0.00156929 |
| | IDH3G | 0.0121836 | 0.0173949 | 0.51372 | 0.030361647 | 2.595608 | 0.009442375 |
| | GLRX3 | 0.02981834 | 0.0339679 | 0.18797 | 0.1968696 | 1.83672 | 0.066251274 |
| | ME1 | 0.02977657 | 0.0227308 | -0.38953 | 0.279251002 | -2.11817 | 0.034160327 |
| | IDH3B | 0.02821311 | 0.0232403 | -0.27973 | 0.321138928 | -2.39981 | 0.016403495 |

| | | | | | | | |
|---|---|---|---|---|---|---|---|
| Olygod | SOD2 | 0.09328127 | 0.0922194 | -0.01652 | 0.356227429 | 0.580785 | 0.56138534 |
| | CAT | 0.04654198 | 0.0390566 | -0.25297 | 0.417613982 | -2.66223 | 0.007762526 |
| | GLRX2 | 0.00941504 | 0.0115999 | 0.30107 | 0.436023935 | 1.098199 | 0.2721178 |
| | PRDX3 | 0.02503786 | 0.0227226 | -0.13998 | 0.465892546 | 0.873362 | 0.38246588 |
| | IDH2 | 0.06061577 | 0.0558009 | -0.11941 | 0.480186804 | -1.67049 | 0.09482261 |
| | GSTK1 | 0.05865564 | 0.0529089 | -0.14876 | 0.51968509 | -1.7879 | 0.073792019 |
| | GPX7 | 0.00177698 | 0.0020373 | 0.19723 | 0.556878645 | | |
| | TXNRD2 | 0.01554967 | 0.0114692 | -0.43912 | 0.576416453 | -1.3224 | 0.186033622 |
| | IDH1 | 0.02677897 | 0.0293803 | 0.13375 | 0.640693732 | 0.220422 | 0.825542184 |
| | ME3 | 0.00550433 | 0.0042631 | -0.36865 | 0.645657278 | | |
| | GPX3 | 0.00285999 | 0.0020743 | -0.4634 | 0.699894949 | | |
| | GSR | 0.0108507 | 0.0118772 | 0.13041 | 0.750870601 | 0.895108 | 0.370729544 |
| | IDH3A | 0.02821681 | 0.0270639 | -0.06019 | 0.77865756 | -0.68324 | 0.494452517 |
| | GPX1 | 0.01813477 | 0.0161061 | -0.17115 | 0.824495792 | -0.99352 | 0.320455658 |
| | GSTM4 | 0.00982193 | 0.0102531 | 0.06198 | 0.89217432 | -0.85299 | 0.393667345 |
| | NNT | 0.05460491 | 0.0525784 | -0.05456 | 0.9060949 | -0.26607 | 0.7901823 |
| | TXN2 | 0.02747294 | 0.0250144 | -0.13525 | 0.9158388 | -0.5944 | 0.5522443 |
| | ME2 | 0.0438445 | 0.0425909 | -0.04185 | 0.9377614 | -0.81992 | 0.4122624 |
| | PRDX5 | 0.04387028 | 0.0411347 | -0.09289 | 0.9674535 | 0.061117 | 0.9512662 |
| | GLRX5 | 0.01041129 | 0.0096397 | -0.11109 | 0.9794574 | -0.38309 | 0.7016501 |
| Microg | GPX1 | 0.11928596 | 0.2128047 | 0.83511 | 0.008253088 | 4.239458 | 2.24E-05 |
| | TXNRD2 | 0.01897567 | 0.0086088 | -1.14027 | 0.29008051 | -1.86761 | 0.061816171 |
| | GPX4 | 0.03715587 | 0.0626959 | 0.75478 | 0.389842164 | 2.056589 | 0.039725758 |
| | PRDX3 | 0.00664249 | 0.0128812 | 0.95548 | 0.393893537 | 0.87784 | 0.380030296 |
| | NNT | 0.04820652 | 0.0677334 | 0.49064 | 0.61568646 | 1.767496 | 0.077145273 |
| | IDH3A | 0.03022823 | 0.0207602 | -0.54208 | 0.624098739 | -2.34172 | 0.019195191 |
| | GSTP1 | 0.0268314 | 0.0385531 | 0.52292 | 0.630370956 | 1.910158 | 0.056112848 |
| | GPX7 | 0.00741648 | 0.0027628 | -1.42459 | 0.644755628 | | |
| | ME3 | 0.01886195 | 0.0083923 | -1.16834 | 0.669456063 | -0.92798 | 0.353419634 |
| | CAT | 0.00936299 | 0.0205904 | 1.13693 | 0.677525015 | 2.075175 | 0.037970352 |
| | PRDX5 | 0.01472771 | 0.0256186 | 0.79866 | 0.720414928 | 1.699241 | 0.089273755 |
| | IDH3G | 0.01128389 | 0.0158667 | 0.49174 | 0.743362335 | 0.555833 | 0.578325364 |
| | GLRX3 | 0.02855771 | 0.0392769 | 0.4598 | 0.751613254 | 1.113826 | 0.265353652 |
| | ME2 | 0.10811705 | 0.1205889 | 0.1575 | 0.775119042 | 0.426205 | 0.669958384 |
| | GSTK1 | 0.03587587 | 0.040067 | 0.1594 | 0.850169478 | 0.626599 | 0.530922328 |
| | TXN2 | 0.01267348 | 0.0193193 | 0.60823 | 0.880059899 | -1.40248 | 0.160772856 |
| | GSR | 0.02471912 | 0.0204164 | -0.2759 | 0.910879178 | -0.30397 | 0.761150115 |
| | SOD2 | 0.08002331 | 0.0721939 | -0.14854 | 0.93720698 | -0.08469 | 0.932508306 |
| | ME1 | 0.01447103 | 0.0193348 | 0.41803 | 0.949971901 | 0.227203 | 0.820265991 |
| | IDH1 | 0.05222991 | 0.0611369 | 0.22717 | 0.965833717 | 0.04454 | 0.964473651 |
| | IDH2 | 0.06422168 | 0.0574509 | -0.16073 | 0.965833717 | -1.10217 | 0.270387527 |
| | IDH3B | 0.01505324 | 0.0198896 | 0.40194 | 0.967898967 | 0.436665 | 0.662354168 |
| | GSTM4 | 0.01311326 | 0.012126 | -0.11293 | 0.968569903 | -0.61583 | 0.538005058 |

**Supplemental Table 1**
